# The importomer is a peroxisomal membrane protein translocase

**DOI:** 10.1101/2020.05.01.072660

**Authors:** James S Martenson, Hanson Tam, Alexander J McQuown, Dvir Reif, Jing Zhou, Vladimir Denic

## Abstract

Three sites of membrane protein biogenesis (the endoplasmic reticulum, mitochondria and chloroplasts) receive unfolded substrates from organelle-specific protein targeting factors, then integrate them using separate translocation channels. Peroxisomes also receive membrane proteins from known targeting factors, but whether a separate translocase is needed for integration remains unknown. Here, using a novel genetic screening strategy, we reveal that the importomer–known for matrix protein import–integrates the peroxisomal membrane protein Pex14. In importomer mutants, Pex14 is arrested in a pre-integrated state on the peroxisome surface. To undergo integration, a Pex14 translocation signal binds the importomer’s substrate receptor Pex5 at a distinct site from matrix proteins. En bloc translocation of an engineered protein complex with Pex14’s luminal region argues that integration occurs without substrate unfolding. Our work shows that the handling of membrane protein targeting and integration by discrete machineries is a fundamental principle shared by diverse membrane protein biogenesis pathways.

## INTRODUCTION

Transmembrane proteins mediate critical processes in all three domains of life, and thus facilitating their journey from the cytosol where they are synthesized into specific cell membranes where they perform their functions is a fundamental requirement for life as we know it. The biogenesis of all membrane proteins occurs in two sequential stages: 1) substrate targeting to a specific membrane followed by 2) integration into and translocation across the membrane of select substrate regions (with the notable exception of single-pass transmembrane proteins lacking significant luminal regions (Anghel et al., 2017; Guna et al., 2018; Wang et al., 2014)). In the case of membrane proteins destined for compartments of the secretory/endocytic system in eukaryotes, targeting is mediated by the cytosolic signal recognition particle and its endoplasmic reticulum (ER)-localized receptor. Substrates are then transferred to the Sec61 translocation channel, which conducts unfolded substrates to enable their membrane integration and translocation (Park and Rapoport, 2012). Similarly, membrane protein biogenesis at mitochondria and chloroplasts is mediated by distinct targeting machineries and translocation channels acting sequentially on unfolded substrates (Schleiff and Becker, 2011; Wiedemann and Pfanner, 2017). The peroxisome—a membrane-bound organelle required for a variety of essential metabolic pathways and reactive oxygen species detoxification—is the final known site of membrane protein biogenesis in eukaryotic cells but while the mechanism of its targeting stage has been elucidated, how substrates then undergo integration and translocation remains poorly understood (Hettema et al., 2014). If this latter mechanism were mediated by a discrete translocase acting downstream of targeting, it would illustrate a unifying mechanistic principle of membrane protein biogenesis across four diverse organelles.

Cell-free studies have provided the strongest evidence that newly-synthesized peroxisomal membrane proteins (PMPs) can be post-translationally delivered to peroxisomes for integration (Diestelkötter and Just, 1993; Fujiki et al., 1984; Fujiki, Yukio et al., 1989; Imanaka, Tsuneo et al., 1996; Pause et al., 1997). This targeting mechanism is mediated by the cytosolic factor Pex19, which can simultaneously bind to its PMP substrates and dock to its peroxisome receptor, the single-pass PMP Pex3 (Fang et al., 2004; Hettema et al., 2000; Jones et al., 2004; Sacksteder et al., 2000). In the case of the tail-anchored PMP Pex26, some evidence argues that Pex3 could itself be an insertase (Chen et al., 2014). However, the question of whether additional machinery mediates integration of PMPs with significant regions on the matrix side has remained unaddressed (Hettema et al., 2014). Complicating this issue is the fact that PMPs can also be first targeted to the ER for integration before being trafficked to peroxisomes in the so-called indirect pathway. The prototype substrate for this pathway is Pex3, which was shown to appear first in the ER when reintroduced into Pex3-deficient cells before appearing in *de novo* generated peroxisomes (Hoepfner et al., 2005). Subsequent studies, including ribosome profiling analyses of ER- and signal recognition particle-associated ribosomes in yeast, argue that multiple PMPs rely on the Sec61 translocation channel for their integration, and that the indirect pathway is a physiological trafficking route *in vivo* (Costa et al., 2018; Jan et al., 2014; van der Zand et al., 2010; van der Zand et al., 2012).

To better define mechanisms by which membrane proteins are directly integrated into peroxisomes, we looked for a model PMP in budding yeast that satisfies two criteria. First, it has a well-defined topology with a matrix-localized domain. Second, it is directly targeted to peroxisomes for integration. We considered the conserved PMP Pex14 as such a model because a rigorous study established that mammalian Pex14 has a single-pass topology with an N-terminal domain (NTD) facing the matrix (Barros-Barbosa et al., 2019). Indeed, we could confirm by *in vitro* protease protection analysis that yeast Pex14 has the same expected topology. We suspected that Pex14 also satisfies the second criterion because an ER-associated ribosome profiling study did not find evidence for its integration by the Sec61 translocation channel (Jan et al., 2014), implying that it is integrated directly into peroxisomes. We could show this was indeed the case by extending the translation time of ribosomes with Pex14 nascent chains and thereby inducing their targeting to peroxisomes as visualized by mRNA fluorescence *in situ* hybridization (FISH).

With Pex14 as our model PMP, we used complementary microscopy and protease protection analyses to identify a region within the Pex14 NTD that is required for translocation of this region during integration but dispensable for Pex14 targeting to peroxisomes. This fortuitous finding implied the existence of a discrete translocase that operated on Pex14 post-targeting. Using a novel Pex14 integration reporter and microscopy, we screened mutants lacking known peroxisomal membrane factors for those that accumulate Pex14 in a pre-integrated state on peroxisomes. Our list of hits was dominated by components of the importomer machinery, which mediates import of soluble matrix proteins in complex with the shuttling factor Pex5. Strikingly, we found that Pex5’s interaction with the Pex14 NTD – independently from its canonical interaction with a peroxisomal targeting signal on its soluble substrates – mediates translocation of this domain in a folded state to facilitate proper Pex14 integration. Lastly, our analysis of Pex14 integration in HEK293T cells argues that the PMP integration mechanism we originally uncovered in yeast is conserved in humans.

## RESULTS

### Yeast Pex14 has an N-terminal matrix domain and is directly targeted to peroxisomes

To validate our first criterion for choosing Pex14 as a model PMP substrate (see Introduction), we first confirmed the N_matrix_-C_cytosol_ topology for yeast Pex14 by performing a protease protection analysis of N-terminally FLAG-tagged and C-terminally MYC-tagged Pex14 (FLAG-Pex14-MYC). In a typical *in vitro* protease protection assay, the membrane protects the protein’s transmembrane and matrix-localized domains from digestion by an exogenous protease. Addition of detergent solubilizes the membrane, thus rendering the membrane-protected fragment sensitive to digestion (Figure 1A). We treated peroxisomes harboring FLAG-Pex14-MYC with two doses of proteinase K and detected a partially-digested intermediate containing the FLAG epitope (~35 kDa) whose decrease was concomitant with an increase in a smaller protected fragment (~18 kDa) corresponding to the expected size for the N-terminal matrix and transmembrane domains. We observed neither species, however, when Triton X-100 was included in the reaction, nor did we see protected fragments containing the C-terminal MYC epitope under any condition tested (Figure 1B). Our results confirm that yeast Pex14 has the expected N_matrix_-C_cytosol_ topology similar to human Pex14 (Barros-Barbosa et al., 2019).

**Figure 1.**
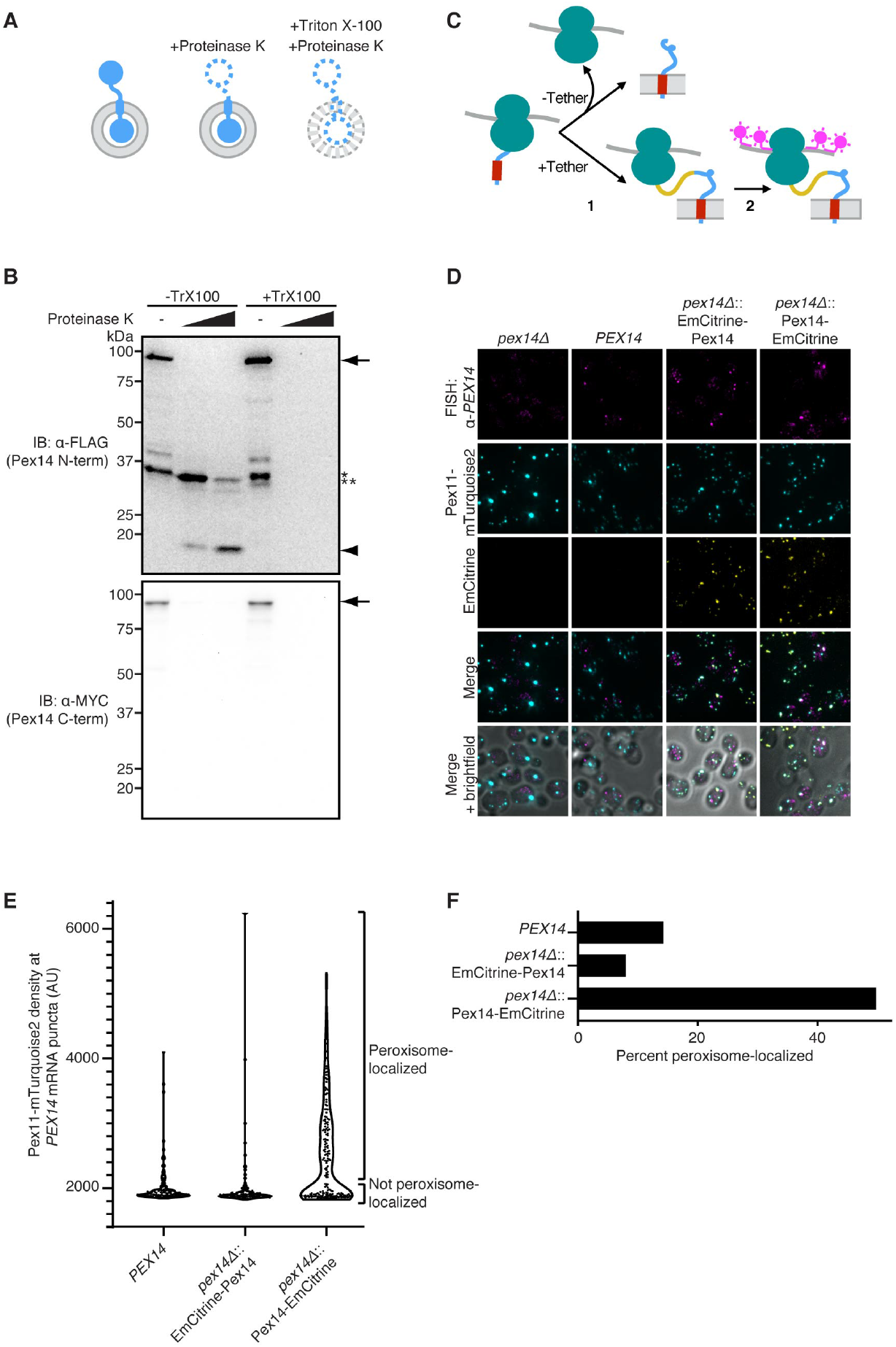
Yeast Pex14 has an N-terminal matrix domain and is directly targeted to peroxisomes. (A) Strategy for defining membrane protein topology using protease protection. (B) Cells expressing FLAG-Pex14-MYC were grown to post-log phase before being subjected to spheroplasting and gentle lysis. Following centrifugation of lysates, membranes were resuspended in buffer with or without Triton X-100 and treated with proteinase K (doses: 0, 31 and 125 ng/μL) for 5 minutes at 37°C as indicated. Following proteinase K quenching, samples were resolved by SDS-PAGE and analyzed by immunoblotting (IB) with the indicated antibodies. Arrow, full length FLAG-Pex14-MYC; arrowhead, protected fragment; asterisk, proteinase K-independent degradation product; double asterisk, proteinase K-dependent partially digested intermediate. (C) Strategy for assessing the direct targeting of Pex14. 1) Post-translationally targeted Pex14 dissociates from its RNC prior to membrane targeting (-Tether), whereas a large C-terminal EmCitrine tag (yellow) tethers the ribosome to the Pex14 nascent chain during membrane targeting. 2) *PEX14* mRNAs are visualized by FISH (FISH probes, magenta). (D) Representative confocal micrographs (maximum intensity projections) of fixed, permeabilized spheroplasts derived from logarithmically growing cells with the indicated *PEX14* locus genotype were stained with anti-*PEX14* mRNA FISH probes. All cells expressed the peroxisomal membrane marker Pex11-mTurquoise2. (E) Distributions of Pex11-mTurquoise2 density at computationally defined *PEX14* mRNA puncta in the images shown representatively in (E). Peroxisome-localized *PEX14* mRNA puncta were defined as having a Pex11-mTurquoise2 density ≥ 2100 arbitrary units (AUs). 147, 125 and 195 *PEX14* mRNA puncta were analyzed for *PEX14, pex14Δ*::EmCitrine-Pex14 and *pex14Δ*::Pex14-EmCitrine strains, respectively. (F) Bar graph of data from (E) showing the fraction of *PEX14* mRNA puncta defined as peroxisomal.

To validate our second criterion, we confirmed the direct targeting of Pex14 to peroxisomes by manipulating the lifetime of Pex14 ribosome-nascent-chain complexes (RNCs) and monitoring any effects this had on mRNA localization using FISH (Figure 1C). While mRNAs of untagged Pex14 or Pex14 with an N-terminal EmCitrine (EmCitrine-Pex14) did not co-localize with the peroxisomal marker Pex11-mTurqouise, a C-terminal EmCitrine fusion (Pex14-EmCitrine) – predicted to extend the lifetime of RNCs with exposed Pex14 signals for peroxisomal targeting – resulted in robust mRNA localization to peroxisomes (Figures 1D-1F). Taken together with the protease protection analysis, these results demonstrate that Pex14 is a suitable model for studying the mechanism of direct PMP integration into peroxisomes.

### The Pex14 NTD carries a translocation signal that is required for integration but dispensable for peroxisomal targeting

We next carried out deletion mutagenesis of a Pex14 tagged with an N-terminal EmCitrine and C-terminal FLAG (EmCitrine-Pex14-FLAG, Figure 2A) to define regions important for peroxisomal targeting. EmCitrine-Pex14-FLAG and its variants expressed at the expected molecular weights and were comparably abundant (Figure 2B). Microscopy analysis revealed that EmCitrine-Pex14-FLAG and a variant lacking the entire Pex14 cytosolic domain (ΔCTD) localized to peroxisomes, whereas the reciprocal variant lacking the N-terminal matrix and transmembrane domains (ΔN171) was mislocalized to the cytosol (Figure 2C). Thus, the N-terminal region containing the matrix and transmembrane domains is both necessary and sufficient for Pex14 targeting to peroxisomes. To further define targeting signals within this region, we deleted the Pex14 transmembrane domain (ΔTMD) and the N-terminal 50 amino acids (ΔN50), which is a region previously implicated in Pex14 targeting (Itoh and Fujiki, 2006; Neufeld et al., 2009). We found that mutants lacking either region alone localized to peroxisomes but a mutant lacking both became mislocalized to the cytosol (Figure 2C). Furthermore, the N-terminal 58 amino acids of Pex14 was sufficient to induce localization of GFP (as N58-GFP) to peroxisomes (Figure 2D). Thus, Pex14 has a distinct N-terminal peroxisomal targeting signal in addition to the targeting information contained within its TMD.

**Figure 2.**
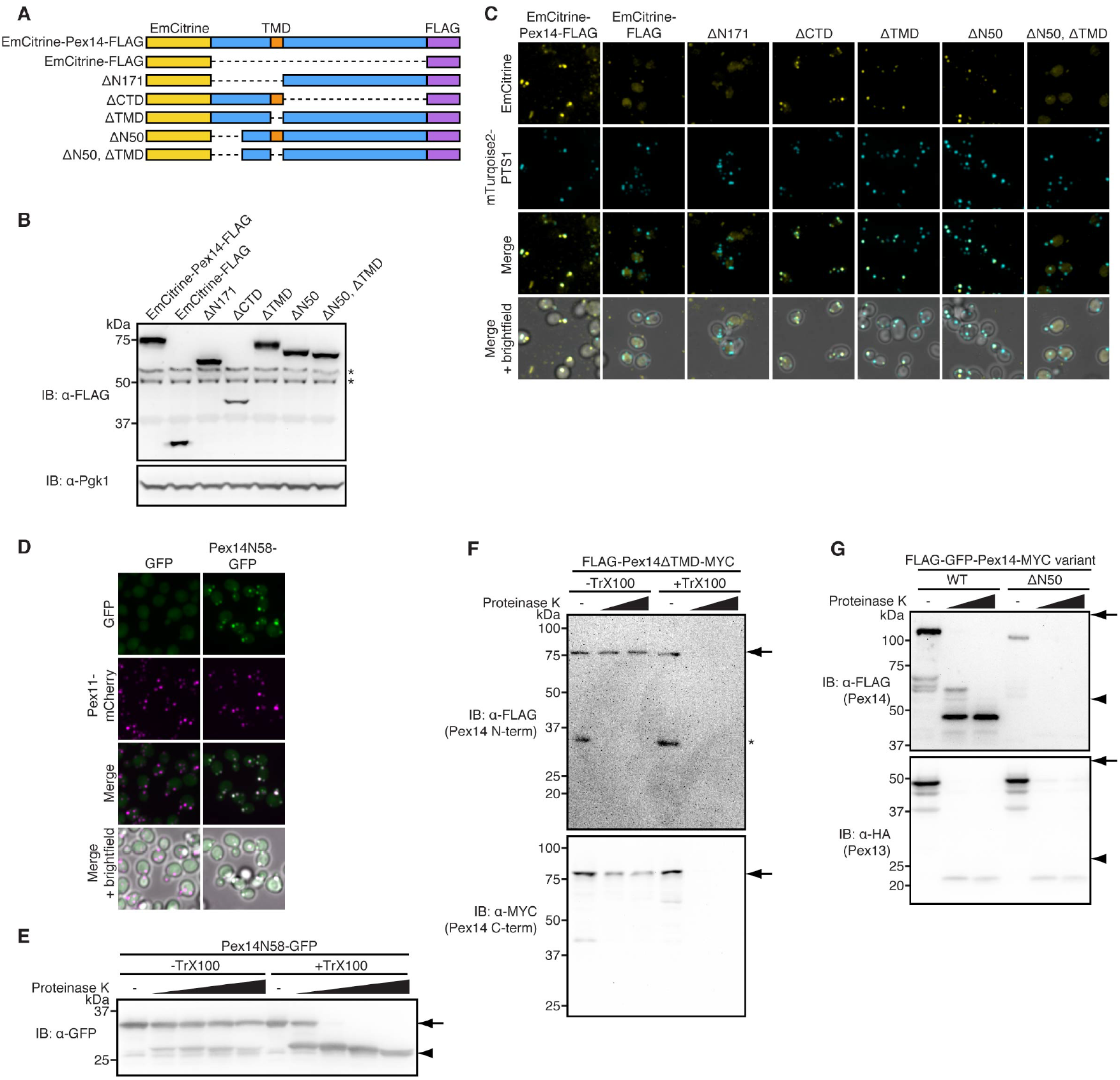
Pex14 carries an N-terminal translocation signal that is dispensable for peroxisomal targeting. (A) Schematic showing EmCitrine- and FLAG-tagged Pex14 and mutants lacking indicated domains or numbered N-terminal amino acid sequences. (B) Extracts from logarithmically-growing cells expressing EmCitrine-Pex14-FLAG or the indicated deletion variants were resolved by SDS-PAGE and analyzed by immunoblotted (IB) with the indicated antibodies. (C) Representative confocal micrographs (maximum intensity projections) of logarithmically growing cells expressing EmCitrine-Pex14-FLAG or the indicated variants and the peroxisomal matrix marker mTurquoise2-PTS1. (D) Representative confocal micrographs (maximum intensity projections) of logarithmically growing cells expressing GFP or Pex14N58-GFP and the peroxisomal membrane marker Pex11-mCherry. (E) Cells expressing Pex14N58-GFP were grown to post-log phase before being subjected to spheroplasting and gentle lysis. Following centrifugation of lysates, membranes were resuspended in buffer with or without Triton X-100, treated with proteinase K (doses: 0, 31, 125, 500 and 2000 ng/μL) for 5 minutes at 37°C. Following proteinase K quenching, samples were resolved by SDS-PAGE and analyzed by immunoblotting (IB) with the indicated antibody. Arrow, full length Pex14N58-GFP; arrowhead, GFP-fragment resistant to proteinase K digestion even in the presence of Triton X-100. (F) Proteinase K protection analysis of membranes derived from cells expressing FLAG-Pex14ΔTMD-MYC was carried out as in (E). Arrow, full length FLAG-Pex14ΔTMD-MYC; asterisk, proteinase K-independent degradation product. (G) Proteinase K protection analysis of membranes derived from cells expressing either FLAG-GFP-Pex14-MYC or FLAG-GFP-Pex14ΔN50-MYC in addition to Pex13-HA was carried out as in (E). Proteinase K doses: 0, 31 and 125 ng/μL. Arrows, full length proteins; arrowheads, fragments resistant to proteinase K digestion.

Why might Pex14 carry a targeting signal within its matrix domain given the robust nature of TMD-dependent Pex14ΔN50 targeting? We hypothesized that this signal enables the matrix domain to undergo translocation across the membrane during Pex14 integration. This model makes the same key prediction for two related constructs: both N58-GFP and Pex14ΔTMD, which both contain the Pex14 N50 targeting signal but lack a predicted TMD, should be fully translocated into the peroxisomal matrix. Indeed, protease protection analysis revealed that the majority of full length N58-GFP (~ 34 kDa) persisted upon addition of increasing amounts of proteinase K, but in the presence of Triton X-100 it yielded an intrinsically proteinase K-resistant GFP fragment (~26 kDa, Figure 2E). Similarly, analysis of FLAG-Pex14ΔTMD-MYC revealed a membrane-dependent protection of its full-length form (as probed for each terminal epitope), and the absence of smaller protected fragments (Figure 2F). Thus, the Pex14 N58 is sufficient for protein translocation into the peroxisome matrix. To test if this region is also necessary for translocation, we analyzed by protease protection full length and N-terminally truncated Pex14-MYC, each carrying a dual FLAG-GFP tag at their N-termini (FLAG-GFP-Pex14-MYC and FLAG-GFP-Pex14ΔN50-MYC, respectively). Consistent with our hypothesis, while FLAG-GFP-Pex14-MYC, upon proteinase K addition, produced the expected sized protected fragment of ~48 kDa corresponding to the FLAG-GFP tag plus the Pex14 matrix and transmembrane domains, we observed no FLAG-tagged protected fragments for Pex14ΔN50 (Figure 2G). As a control, we also examined C-terminally HA-tagged Pex13 (Pex13-HA), a PMP that has been shown to reach peroxisomes indirectly via the ER (Jan et al., 2014; van der Zand et al., 2010), and whose topology has also been analyzed by protease protection. Consistent with published results on mammalian Pex13 (Barros-Barbosa et al., 2019), we found that Pex13-HA’s C-terminus was protected from proteinase K, irrespective of which variant of Pex14 was expressed (Figure 2G). Collectively, these data suggest that the Pex14 NTD instructs its own translocation during Pex14 integration into the peroxisomal membrane.

### The peroxisomal importomer mediates Pex14 integration

The observation that Pex14 variants lacking their first 50 amino acids reach peroxisomes but arrest in a pre-integrated state suggests that a discrete translocation machinery engages the missing N-terminal region as part of the native Pex14 integration pathway. In principle, mutations that ablate such a machinery would enable full-length Pex14 to phenocopy this arrested state. To facilitate genetic screening for potential mutants of this kind, we restricted our search to a collection of gene deletion strains lacking individual peroxins (peroxisome biogenesis factors encoded by the *PEX* genes), as well as other known and predicted PMPs (Yofe et al., 2016). In addition, we devised a facile *in vivo* protease protection assay in which we express the human rhinovirus (HRV) 3C protease in the cytosol and engineer an HRV 3C protease cleavage site into the matrix domain of a C-terminally FLAG-tagged Pex14 reporter (Pex14(hrv)-FLAG, Figure 3A). The extent of cleavage was measured by SDS-PAGE and Western blotting. In the wild-type, we observed only a relatively small fraction of cleaved Pex14 (Figures 2B and S1A), which is likely due to protease accessibility *en route* to reporter integration. Similarly, mutations in a variety of other PMPs involved in peroxisome fission, inheritance, and membrane transport did not significantly affect Pex14 integration as measured by our reporter. Our screen identified Pex3 and Pex19 as hits, which is consistent with their known role in direct targeting of PMPs (see Discussion). More surprisingly, we found strong hits in Pex1, Pex2, Pex4, Pex5, Pex6, Pex8, Pex10, Pex12, Pex13, Pex15, Pex17 and Pex22. Along with Pex14 itself, these factors comprise the well-studied peroxisomal matrix importomer responsible for the import of soluble proteins into the peroxisomal matrix (Hettema et al., 2014). That Pex14 did not turn up as a hit is not surprising, given that the Pex14(hrv)-FLAG reporter itself is functional and complements the *pex14Δ* import defect (data not shown).

**Figure 3.**
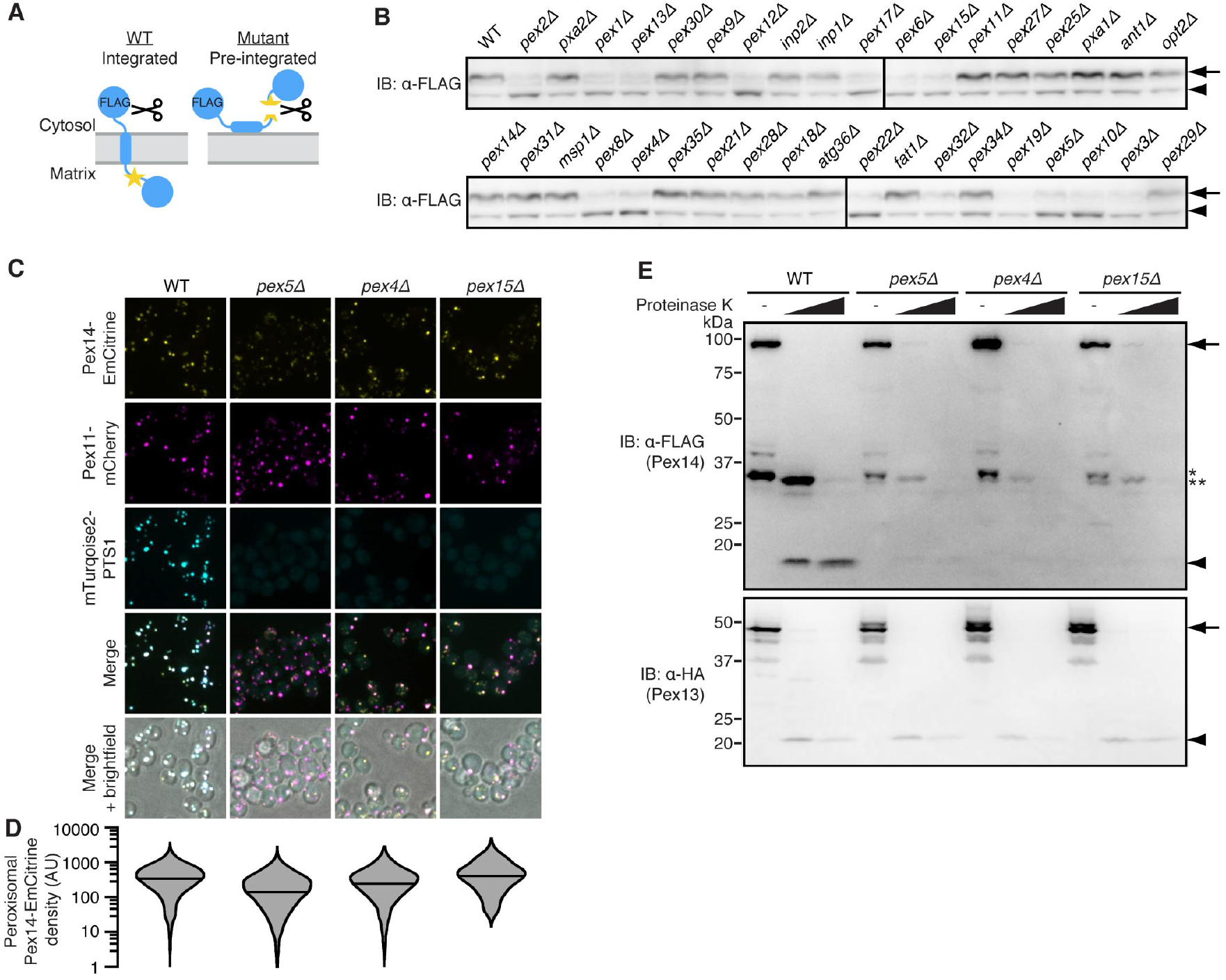
Pex14 becomes arrested in a pre-integrated state on peroxisomes in importomer mutants. (A) Schematic illustrating the strategy for detecting an integration defective mutant using a novel *in vivo* protease assay. The site-specific HRV 3C protease (scissors) is expressed in the cytosol and its cleavage site (star) is engineered into the matrix domain of Pex14. When Pex14 is integrated, the cleavage site is sequestered from the protease. When Pex14 is pre-integrated, the cleavage site becomes accessible to cleavage by the protease. (B) Wild-type (WT) cells or the indicated gene deletions expressing Pex14(hrv)-FLAG under the *PEX14* promoter from the *URA3* locus and cytosolic HRV 3C protease under the *GAL1* promoter were grown logarithmically for 15 hours in 2%/2% galactose/raffinose followed by 2 hours in 2% glucose. Extracts were resolved by SDS-PAGE and analyzed by immunoblotting (IB) with an anti-FLAG antibody. Arrow, full length Pex14(hrv)-FLAG; arrowhead, cleaved Pex14(hrv)-FLAG. (C) Representative confocal micrographs (maximum intensity projections) of logarithmically growing cells expressing Pex14-EmCitrine and the peroxisomal membrane and matrix markers Pex11-mCherry and the mTurquoise2-PTS1, respectively. (D) Distributions of Pex14-EmCitrine density at Pex11-mCherry-defined peroxisomes in the images shown representatively in (C). Medians are represented as horizontal lines. Violin plot tails were cut off at 1 arbitrary units (AUs) for clarity of presentation. 2865, 3204, 2703 and 808 peroxisomes were analyzed for WT, *pex5Δ, pex4Δ* and *pex15Δ* strains, respectively. (E) Cells expressing FLAG-Pex14-MYC and Pex13-HA were grown to post-log phase before being subjected to spheroplasting and gentle lysis. Following centrifugation of lysates, membranes were resuspended in buffer and treated with proteinase K (doses: 0, 31 and 125 ng/μL) for 5 minutes at 37°C as indicated. Following proteinase K quenching, samples were resolved by SDS-PAGE and analyzed by immunoblotting (IB) with the indicated antibodies. Arrow, full length proteins; arrowhead, protected fragments; asterisk, proteinase K-independent degradation product; double asterisk, proteinase K-dependent partially digested intermediate.

Two additional lines of evidence argue that importomer mutations cause Pex14 to phenocopy the pre-integrated state of Pex14ΔN50. First, we measured Pex14 (tagged with a C-terminal EmCitrine) abundance at peroxisomes by quantitative cell microscopy (expressed as EmCitrine fluorescence density at Pex11-mCherry-defined peroxisomes). This analysis revealed that Pex14 targeting largely persisted, albeit to a variable degree, in the importomer mutants we tested (Figures 3C, 3D, S1B and S1C). Second, we complemented this analysis by also analyzing the integration status of FLAG-Pex14-MYC in the corresponding importomer mutants using the *in vitro* protease protection assay. In wild-type cells we observed the ~18 kDa protected fragment indicative of Pex14 integration, whereas this fragment was absent in all the mutants (Figures 3E and S1D). Importantly, we found that the C-terminus of Pex13 remained protected in the mutants arguing that the peroxisomal membrane has not been grossly compromised. Taken together, these data suggest that the importomer plays a direct role in Pex14 integration into the peroxisomal membrane.

### Pex5 binding to the Pex14 NTD enables Pex14 integration independently of PTS1 import

We next turned our focus to the mechanism by which the importomer could mediate Pex14 integration. In its role in matrix protein import, the shuttle factor Pex5 recognizes the type I peroxisomal targeting signal (PTS1) of newly-synthesized matrix proteins to deliver them to the peroxisome surface by binding the docking subcomplex (Pex13, Pex14, Pex17). In a poorly defined step, Pex5 transiently visits the peroxisomal matrix to deposit its cargo. It is then monoubiquitinated by the E2 (Pex4, Pex22) and E3 (Pex2, Pex10, Pex12) subcomplexes, the latter of which is bridged to the docking complex by Pex8. Lastly, the AAA+ transmembrane complex (Pex1, Pex6, Pex15) dislocates monoubiquitinated Pex5 to the cytosol for deubiquitination and further rounds of targeting and import (Hettema et al., 2014).

Pex14 is known to have two Pex5 binding sites. One site, localized to the Pex14 cytosolic C-terminus, is thought to serve as an initial binding site for cargo-laden Pex5 on the docking complex. The other site lies within the evolutionarily conserved region in the Pex14 matrix-localized N50 (Figure 4A) (Niederhoff et al., 2005), and its interaction with Pex5 has been characterized by NMR at atomic resolution (Figure 4B) (Neufeld et al., 2009). Indeed, we observed binding between an *in vitro* synthesized Pex14 variant lacking the C-terminal Pex5 binding domain (ΔC) and recombinant FLAG-tagged Pex5 (FLAG-Pex5). This interaction was specific because it could be disrupted by a mutation in a highly conserved phenylalanine, which makes key pi-stacking interactions with Pex5 (F19A, ΔC; Figures 4C and S2) (Neufeld et al., 2009). We postulated that Pex5 binding to the Pex14 NTD enables the Pex14 NTD to be translocated across the peroxisomal membrane during Pex14 integration. Consistent with this idea, we found that EmCitrine-Pex14ΔTMD was dependent upon Pex5 for its peroxisomal localization, whereas EmCitrine-Pex14 and EmCitrine-Pex14ΔN50 were not (Figure 4D). Immunoblotting analysis revealed that the expression level of mislocalized EmCitrine-Pex14ΔTMD (or the two control constructs) was unaffected by the absence of Pex5 (Figure 4E). Moreover, a Pex14 variant harboring the F19A mutation that disrupts N50 binding to Pex5 failed to integrate into peroxisomes as assessed by protease protection (Figure 4F). These data suggest that Pex5 facilitates translocation of the Pex14 NTD across the membrane during Pex14 integration.

**Figure 4.**
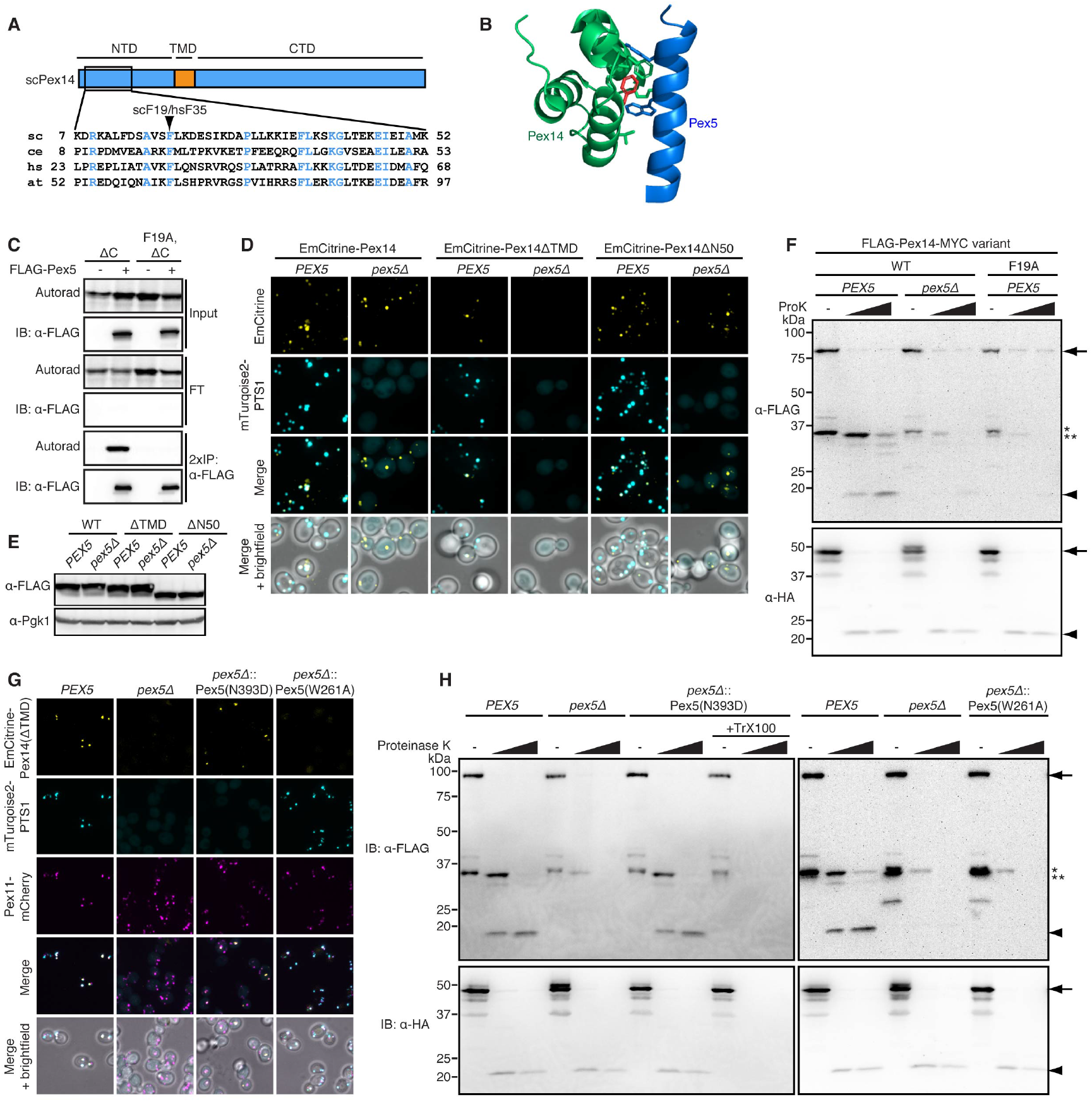
Pex5 mediates Pex14 integration independently of PTS1 import by binding to the Pex14 NTD. (A) (above) Schematic depicting Pex14’s domain organization. NTD: N-terminal domain, TMD: transmembrane domain, CTD: C-terminal domain. (below) Amino acid alignment of the indicated NTD region from the following Pex14 homologs: Saccharomyces cerevisiae (sc), Caenorhabditis elegans (ce), Homo sapiens (hs) and Arabidopsis thaliana (at). Absolutely conserved residues are highlighted in blue. (B) Solution NMR structure of the conserved region of the human Pex14 NTD bound to a Pex5-derived peptide (PDB2w84, Neufeld et al., 2009). The key interacting phenylalanine F35 (corresponding to F19 in yeast) on Pex14 is highlighted. (C) *In vitro* translation of mRNAs encoding the indicated Pex14 variants in the presence of S35-labeled methionine and recombinant FLAG-Pex5 was followed by anti-FLAG immunoprecipitation. Input, flowthrough (FT), and 2x-loaded immunoprecipitation (2xIP) samples were resolved by SDS-PAGE and analyzed by autoradiography (Autorad) and immunoblotting (IB) with anti-FLAG. (D) Representative confocal micrographs (maximum intensity projections) of logarithmically growing cells of the indicated genotypes expressing EmCitrine-Pex14-FLAG variants and the peroxisomal matrix marker mTurquoise2-PTS1. (E) Extracts from logarithmically-growing cells of the indicated genotypes expressing EmCitrine-Pex14-FLAG or the indicated deletion variants were resolved by SDS-PAGE and analyzed by immunoblotting with the indicated antibodies. (F) Cells expressing the indicated FLAG-Pex14-MYC variant (WT or F19A) and Pex13-HA were grown to post-log phase before being subjected to spheroplasting and gentle lysis. Following centrifugation of lysates, membranes were resuspended in buffer and treated with proteinase K (doses: 0, 31 and 125 ng/μL) for 5 minutes at 37°C as indicated. Following proteinase K quenching, samples were resolved by SDS-PAGE and analyzed by immunoblotting (IB) with the indicated antibodies. Arrow, full length proteins; arrowhead, protected fragments; asterisk, proteinase K-independent degradation product; double asterisk, proteinase K-dependent partially digested intermediate. (G) As in (D), except cells had the indicated *PEX5* locus genotypes and also expressed the peroxisomal membrane marker Pex11-mCherry. (H) Proteinase K protection analysis of membranes derived from cells with the indicated *PEX5* locus genotypes expressing FLAG-Pex14-MYC and Pex13-HA was carried out with or without Triton X-100 as in (F). Arrow, full length proteins; arrowhead, protected fragments; asterisk, proteinase K-independent degradation product; double asterisk, proteinase K-dependent partially digested intermediate.

To further substantiate this model, we next defined mutations in Pex5 that functionally separated its role in Pex14 integration from its role in import of PTS1-containing cargo. We focused on two mutants: Pex5(N393D), which disrupts PTS1 binding (Klein et al., 2002), and Pex5(W261A), which disrupts a distinct site involved in binding to the Pex14 N50 (Kerssen et al., 2006). We found that Pex5(N393D) enabled accumulation of EmCitrine-Pex14ΔTMD, but not the peroxisomal matrix marker mTurquoise2-PTS1, at peroxisomes. The converse was true for Pex5(W261A): It enabled peroxisomal accumulation of mTurquoise2-PTS1 but not EmCitrine-Pex14ΔTMD (Figure 4G). Protease protection analysis of these mutants revealed that Pex14 integration was selectively disrupted in Pex5(W261A) but not Pex5(N393D) (Figure 4H). Collectively, our dissection of Pex5-cargo interactions strongly supports a model in which the importomer mediates Pex14 integration across the peroxisomal membrane independently of PTS1 import.

### The Pex5-Pex14 NTD integration pathway translocates a folded protein complex

To further define how Pex5 facilitates Pex14 integration, we considered two mechanistic models. The Pex14 NTD might be translocated in a folded state, consistent with the well documented ability of the importomer to translocate folded PTS1 substrates (Glover et al., 1994; McNew and Goodman, 1994; Romano et al., 2019; Walton et al., 1995) and the observation that the Pex5-bound Pex14 NTD is folded (Figure 4B) (Neufeld et al., 2009). Alternatively, the Pex14 NTD might be translocated in an unfolded state in a manner akin to the translocation channel-mediated translocation at other organelles. To distinguish between these possibilities, we asked if an otherwise cytosolic protein lacking PTSs becomes imported into peroxisomes if non-covalently tethered to the Pex5-Pex14 NTD complex. For our piggyback strategy, we fused one half of a previously characterized coiled-coil to the Pex14 N58 (CC(-)-FLAG-Pex14N58) and the other (complementary) half of the coiled-coil to GFP (CC(+)-GFP, Figure 5A) (Moll et al., 2001). We found that, when expressed alone under the strong *TDH3* promoter, CC(+)-GFP localized diffusely in the cytosol, but when co-expressed with CC(-)-FLAG-Pex14N58 (also under the *TDH3* promoter), it accumulated at peroxisomes (Figure 5B) where a substantial fraction of full length CC(+)-GFP was protease resistant in the absence, but not presence, of Triton X-100 (Figure 5C). These results suggest that the importomer translocates not only soluble proteins but also luminal regions of membrane proteins in their folded state.

**Figure 5.**
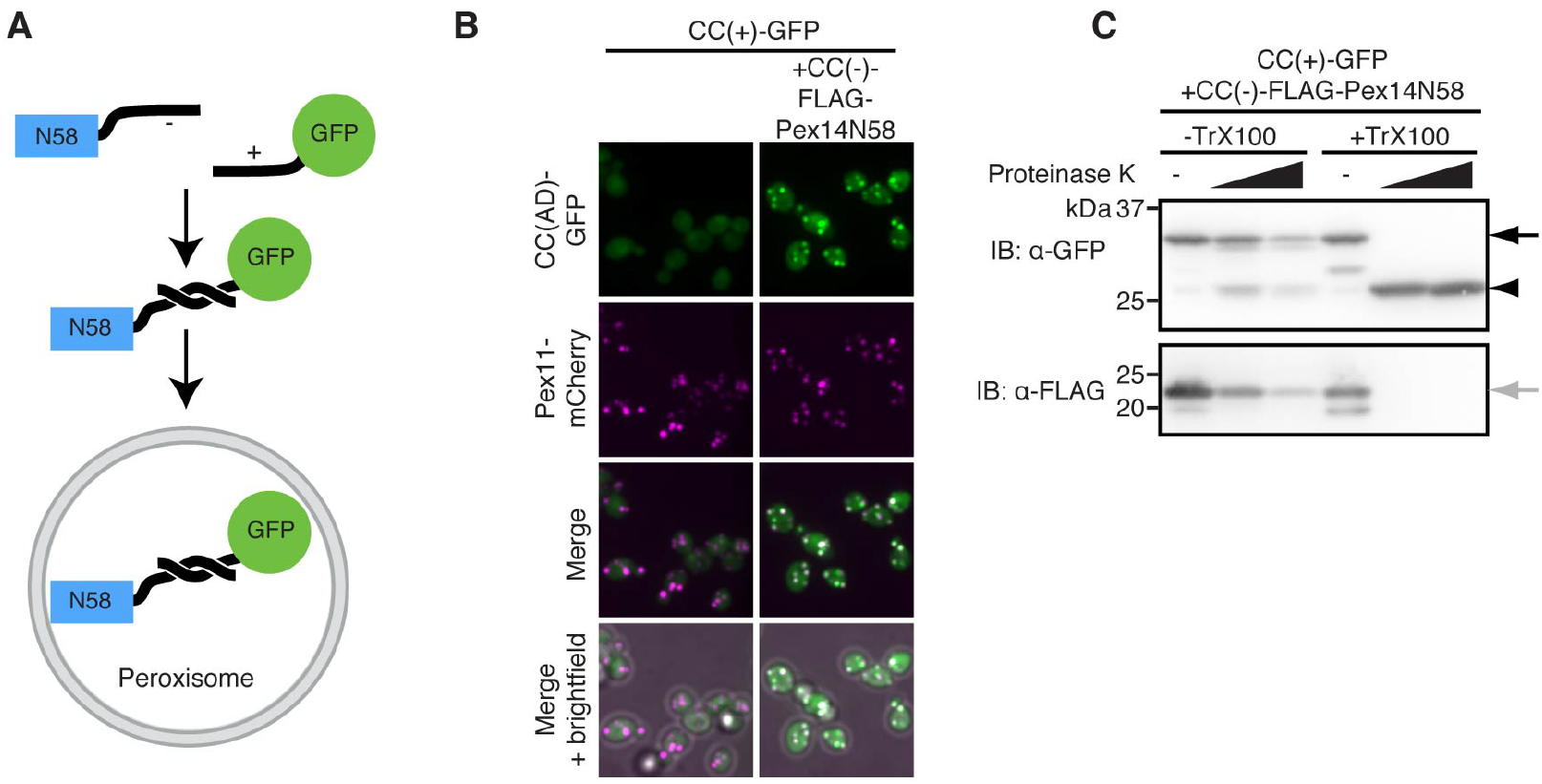
Translocation of the Pex14 NTD without unfolding. (A) “Piggy-backing” strategy for testing if Pex14 NTD can undergo membrane translocation in a folded state. One half of a coiled-coil (-) is fused to the Pex14 N58 and the other half (+) to GFP. The appearance of GFP in the matrix implies that it underwent peroxisomal targeting and translocation as part of a coiled-coil complex. (B) Representative confocal micrographs (maximum intensity projections) of logarithmically growing cells expressing CC(+)-GFP and CC(-)-FLAG-Pex14N58 as indicated under *TDH3* promoters. Cells also expressed the peroxisomal membrane marker Pex11-mCherry. (C) Cells expressing CC(+)-GFP and CC(-)-FLAG-Pex14N58 were grown to post-log phase before being subjected to spheroplasting and gentle lysis. Following centrifugation of lysates, membranes were resuspended in buffer with or without Triton X-100, treated with proteinase K (doses: 0, 125 and 500 ng/μL) for 5 minutes at 37°C. Following proteinase K quenching, samples were resolved by SDS-PAGE and analyzed by immunoblotting (IB) with the indicated antibodies. Arrow, full length proteins; arrowhead, GFP-fragment resistant to proteinase K digestion even in the presence of Triton X-100.

### Evidence that importomer-mediated Pex14 integration is conserved in humans

The fact that residues in the Pex14 NTD involved in Pex5 binding are conserved from yeast to humans suggests that importomer-mediated Pex14 integration is also conserved in human cells. To test this idea, we generated Pex5 knock out (*PEX5^KO^*) HEK293T cells using CRISPR/Cas9 technology, and assessed the effect on Pex14 peroxisomal targeting and integration. Sequencing of the *PEX5* locus confirmed chromosomal disruptions leading to premature stop codons in both alleles (Figures S3A-S3C). We further confirmed by immunoblotting that Pex5 was absent in *PEX5^KO^* cells but that this ablation had only minimal effects on the expression of endogenous Pex14 and N-terminally FLAG and C-terminally HA tagged Pex26 (FLAG-Pex26-HA, introduced via lentiviral transduction) (Figure 6A). Pex14 and FLAG-Pex26-HA both colocalized with the peroxisomal marker PMP70 in WT and *PEX5^KO^* cells, whereas the peroxisomal matrix protein catalase, which relies on the importomer for its import, became diffusely cytosolic in the absence of Pex5 (Figure 6B). Consistent with previously published results, our protease protection analysis with a Pex14 antibody known to recognize the Pex14 matrix/transmembrane domains (Barros-Barbosa et al., 2019) revealed a protected, Triton X-100-sensitive fragment (~18 kDa) corresponding to these domains (Figure 6C). This fragment was absent in *PEX5^KO^* cells, even after adjusting loading to match input levels from WT cells. As a control for membrane integrity, we established that the expected membrane protected fragment of FLAG-Pex26-HA persisted in *PEX5^KO^* cells. These results argue that in human cells, similar to yeast, the importomer is required for a discrete Pex14 NTD translocation step that can be uncoupled from Pex14 peroxisomal targeting.

**Figure 6.**
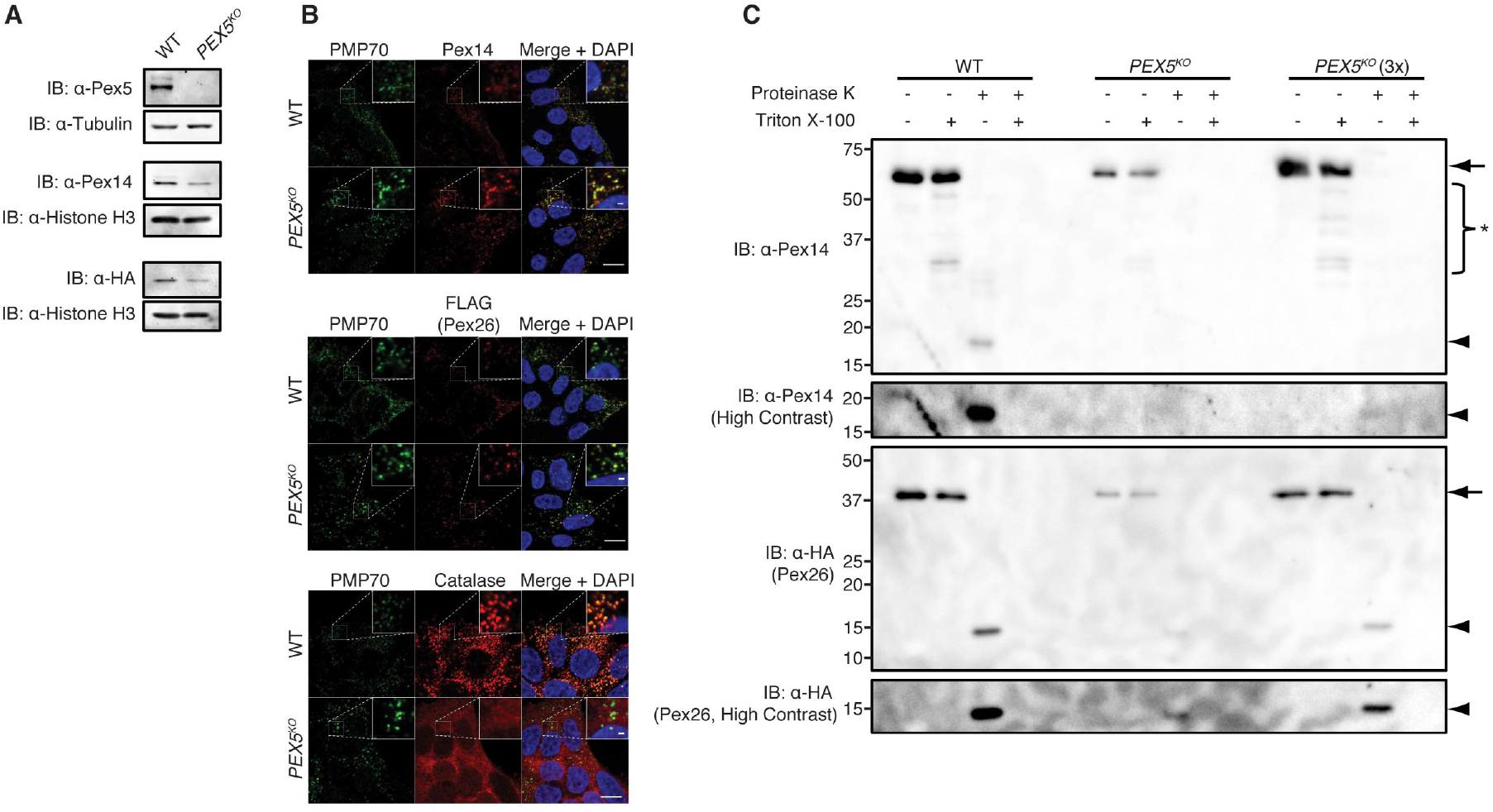
Evidence for conservation of importomer-mediated Pex14 integration in humans. (A) Extracts from HEK293T cells of the indicated genotypes expressing FLAG-Pex26-HA (introduced via lentiviral transduction and expressed under control of the UbC promoter) were resolved by SDS-PAGE and analyzed by immunoblotting (IB) with the indicated antibodies. (B) Representative confocal micrographs (single Z plane near the center of the cell) of fixed, permeabilized HEK293T cells of the indicated genotypes expressing FLAG-Pex26-HA (introduced via lentiviral transduction and expressed under control of the UbC promoter) and stained with anti-PMP70, anti-Pex14, anti-FLAG and anti-calatase antibodies and DAPI as indicated. Endogenous Pex14 and exogenous FLAG-Pex26-HA colocalized with the peroxisomal marker PMP70 in Pex5 WT and KO cells, whereas catalase colocalized with PMP70 in Pex5 WT cells but became diffusely cytosolic in *PEX5^KO^* cells. (C) HEK293T of the indicated *PEX5* genotype expressing endogenous Pex14 and exogenous FLAG-Pex26-HA (introduced via lentiviral transduction and expressed under control of the UbC promoter) were subjected to gentle lysis. Following centrifugation of lysates, membranes were resuspended in buffer with or without Triton X-100 and treated with proteinase K (doses: 0 and 200 ng/μL) for 40 minutes on ice. Following proteinase K quenching, samples were resolved by SDS-PAGE and analyzed by immunoblotting (IB) with the indicated antibodies. Arrows, full length proteins; arrowheads, protected fragments; asterisk, partial cleavage products.

## DISCUSSION

With the exception of TA protein insertases, machines for integrating transmembrane helices at the ER, mitochondria and chloroplasts as well as the prokaryotic plasma membrane rely on narrow translocation channels that conduct unfolded protein regions across lipid bilayers (Anghel et al., 2017; Guna et al., 2018; Park and Rapoport, 2012; Schleiff and Becker, 2011; Wang et al., 2014; Wiedemann and Pfanner, 2017), but whether PMP targeting to the peroxisome is followed by a mechanistically analogous strategy for entering the membrane has remained unknown. To probe this issue, we focused our study on the topological state of Pex14 at the peroxisome membrane. This choice for a PMP model substrate was informed by our fortuitous discovery that a Pex14 lacking an N-terminal sequence was still be targeted to the peroxisome surface only to stall along its biogenesis pathway in a pre-integrated state. We then demonstrated that Pex14’s integration can also be separated from its targeting at the level of the machineries mediating these two distinct stages of its biogenesis. Specifically, in importomer mutants, we found that the vast majority of Pex14 remains correctly targeted to the peroxisome but not integrated. To achieve Pex14 integration, the importomer uses a mechanism akin to its canonical ability to translocate fully folded soluble proteins across the peroxisome membrane. To our knowledge this is the first example of a membrane protein translocase capable of translocating a luminal membrane protein region in a folded state.

Our work raises important questions about the dynamics of importomer-mediated PMP integration. For example, how is it that a highly dynamic pore known to conduct a wide size range of folded soluble substrates (Glover et al., 1994; McNew and Goodman, 1994; Meinecke et al., 2010; Romano et al., 2019; Walton et al., 1995) can terminate further protein translocation upon encountering the Pex14 TMD? Is the Pex14 TMD partitioned laterally into the bilayer by dissolution of a dynamic transmembrane pore? How might this be linked to Pex5 dislocation by the AAA+ transmembrane complex (Platta et al., 2005)? The *in vitro* reconstitution of importomer-mediated PMP integration into proteoliposomes using purified components would provide a critical tool for dissecting these mechanistic questions.

Another important question concerns the degree to which other directly targeted PMPs share Pex14’s importomer-mediated integration pathway. A key importomer component is the shuttle factor Pex5, which in its canonical role binds the PTS1 of many matrix proteins via its tetratricopeptide repeat domain (Brocard et al., 1994; Leij et al., 1993; McCollum et al., 1993). To mediate Pex14 integration Pex5 uses a distinct, noncanonical site to bind to and translocate Pex14’s NTD. This noncanonical site was also shown to mediate Pex5-dependent, PTS1-independent import of the peroxisomal matrix proteins Pox1 and Cat2 (Klein et al., 2002), indicating that they share the translocation pathway utilized by Pex14. More generally, systematic efforts to define additional proteins that bind this noncanonical site in Pex5 could reveal the extent to which other PMPs and matrix proteins share Pex14’s translocation strategy. Our results do not preclude, however, the possibility that additional PMP integration pathways exist, especially in the case of PMPs with complex membrane topologies such as those assumed by several peroxisomal ABC transporters. The HRV 3C protease screening platform used in our study is a proof-of-concept tool for further exploring the diversity of peroxisomal integration pathways using other PMPs as substrates.

Our findings also provide insight into the mechanism of importomer-mediated matrix protein import. It has been speculated that Pex5 binding to Pex14’s NTD is a key step in the PTS1 import cycle (Emmanouilidis et al., 2016). In light of this, we considered the possibility that Pex14 integration was concomitant with PTS1 protein import but instead found, consistent with a previous study, that a Pex5 mutant unable to bind Pex14 (W261A) still mediated PTS1 import (Figure 4G) (Kerssen et al., 2006). While Pex14 is not appreciably integrated in this mutant background (Figure 4H), we can’t exclude the possible existence of a small fraction of integrated Pex14 (undetectable by our protease protection assay) that is sufficient to catalytically import matrix proteins under cell growth conditions that don’t necessitate peroxisome function. Regardless, Pex5-mediated Pex14 integration does not appear to be fundamental to the PTS1 import cycle.

While the importomer factors represented the most interesting hits from our HRV 3C protease screen, several other genes on our list warrant further commentary. First, our screen identified the canonical PMP targeting factor Pex19 and its membrane anchor Pex3 as essential for protecting our Pex14 reporter from HRV 3C protease cleavage (Figure 3B). In Pex19 and Pex3 null cells, mature peroxisomes are absent. Instead, a subset of PMPs including Pex14 and some, but not all, of the other importomer subunits, localize to small peroxisomal remnants, which may serve as precursors for peroxisome generation (Knoops et al., 2014; Wróblewska et al., 2017). Consistent with the incomplete compliment of importomer subunits at these structures, our screen results imply that Pex14 might be in a pre-integrated state on these peroxisomal remnants, but further analysis is needed to definitively test Pex14’s integration status on these structures. Another unresolved question is whether, beyond its role in PMP targeting, Pex3 also functions directly in PMP integration into lipid bilayers. One piece of evidence supporting this model is the observation that a mutant of *Neurospora* Pex3 that interacts normally with Pex19 nonetheless fails to insert the tail-anchored protein Pex26, suggesting that this mutant may have a defect in Pex26 insertion specifically (Chen et al., 2014). However, beyond the special case of tail-anchored proteins, no Pex3 separation of function mutant that is able to target PMPs to peroxisomes but leaves them stalled in a pre-integrated state has been reported. Lastly, we noted the absence from our screen hits of type 2 peroxisomal targeting signal (PTS2)-specific import factors Pex7, Pex18 and Pex21 (Hettema et al., 2014), suggesting that they engage the importomer independently from the Pex14 integration pathway.

In addition to membrane integration, our results also shed light on Pex14 targeting. We found that extending the lifetime of Pex14 RNCs with exposed peroxisomal targeting signals via a C-terminal EmCitrine “tether” caused them to localize at peroxisomes (Figures 1D-1F). The result suggests not only that Pex14 is targeted directly to peroxisomes, but that direct Pex14 targeting under standard nutrient conditions is primarily post-translational. (A limitation of this result is that it does not indicate the proportion of directly versus indirectly targeted Pex14. However, the absence of integrated Pex14 in importomer mutants suggests a minimal contribution from the latter targeting mechanism.) Interestingly, a previous study reported a similar redistribution of *Pex14* mRNA from the cytosol to peroxisomes when cells were shifted from standard media to media containing oleate as the carbon source (a condition under which peroxisomes perform a vital catabolic function) (Zipor et al., 2009). As a parsimonious synthesis with our own finding, we speculate that this shift is due to a general reduction in the rate of RNC elongation under suboptimal growth conditions. Regardless, the prevalence of co-versus post-translational direct PMP targeting could be assessed systematically using ribosome profiling to analyze peroxisome-associated RNCs in both standard and oleate media and identify new oleate-inducible factors that enhance the kinetics of Pex14 targeting. We were also surprised to identify two independent peroxisomal targeting signals within Pex14 – the N50 and TMD. While we demonstrated that the N50 relies on Pex5 for targeting, the targeting factor that engages the TMD remains unknown. The simplest possibility is that the Pex14 TMD utilizes the canonical PMP targeting factor Pex19, although future studies will have to test this idea formally.

Finally, a controversy in the field of peroxisome biogenesis has been whether peroxisomes form exclusively *de novo* from the ER, or whether they multiply by growth and division (Hettema et al., 2014; Hoepfner et al., 2005; Motley et al., 2015; Motley and Hettema, 2007; van der Zand et al., 2010; van der Zand et al., 2012). Implicit in the growth and division model is the idea that most PMPs do not rely on the ER-localized Sec61 translocation channel for their integration, but instead are directly targeted and integrated into peroxisome membranes. Despite a wealth of evidence supporting direct PMP targeting by the targeting factors Pex3 and Pex19, a missing piece has been the identity of a peroxisome-localized PMP translocase (Fang et al., 2004; Hettema et al., 2014, 2000; Jones et al., 2004; Liu et al., 2016; Sacksteder et al., 2000). Here we identify the importomer as a PMP translocase, and thus contribute to the accumulating evidence supporting the growth and division model.

To conclude, our work reveals the versatility of the importomer as a translocation machine not only for soluble but also for membrane proteins residing in the peroxisome, both of which can be translocated in a folded state. At the broader level of membrane biology, our work indicates that, like the ER, mitochondria, chloroplasts and the bacterial plasma membrane, peroxisomes utilize separate machineries for the targeting and integration of membrane proteins.

## ACKNOWLEDGEMENTS

We thank the Harvard Center for Biological Imaging for infrastructure and support; the Bauer Core Facility at Harvard University for expert technical assistance; Jorge Azevedo for the anti-Pex14 antibody; Didier Trono for plasmid psPAX2; Bob Weinberg for plasmid pCMV-VSV-G; Shankar Mukherji for experimental assistance with RNA FISH; Nick Weir and Vivian Jou for experimental assistance with the protease protection assays; Houqing Yu for experimental assistance with protein purification; Charlene Chan for experimental assistance with *in vitro* translation; Jeff Offermann for general technical support; Peter Arvidson for administrative assistance; Roarke Kamber for critical reading of the manuscript and members of the Denic lab for helpful scientific discussion. A.J.M. was supported by the National Science Foundation Graduate Research Fellowship under Grant No. DGE1745303.

## AUTHOR CONTRIBUTIONS

Conceptualization, J.S.M. and V.D.; Methodology, J.S.M. and A.J.M.; Formal Analysis, J.S.M. and H.T.; Investigation, J.S.M., H.T., A.J.M. and D.R.; Resources, J.S.M., H.T., A.J.M., D.R. and J.Z.; Writing – Original Draft, J.S.M. and V.D.; Writing – Review & Editing, J.S.M., H.T., A.J.M., D.R. and V.D.; Funding Acquisition, A.J.M. and V.D.; Supervision, J.S.M. and V.D.

---

**Figure S1.**
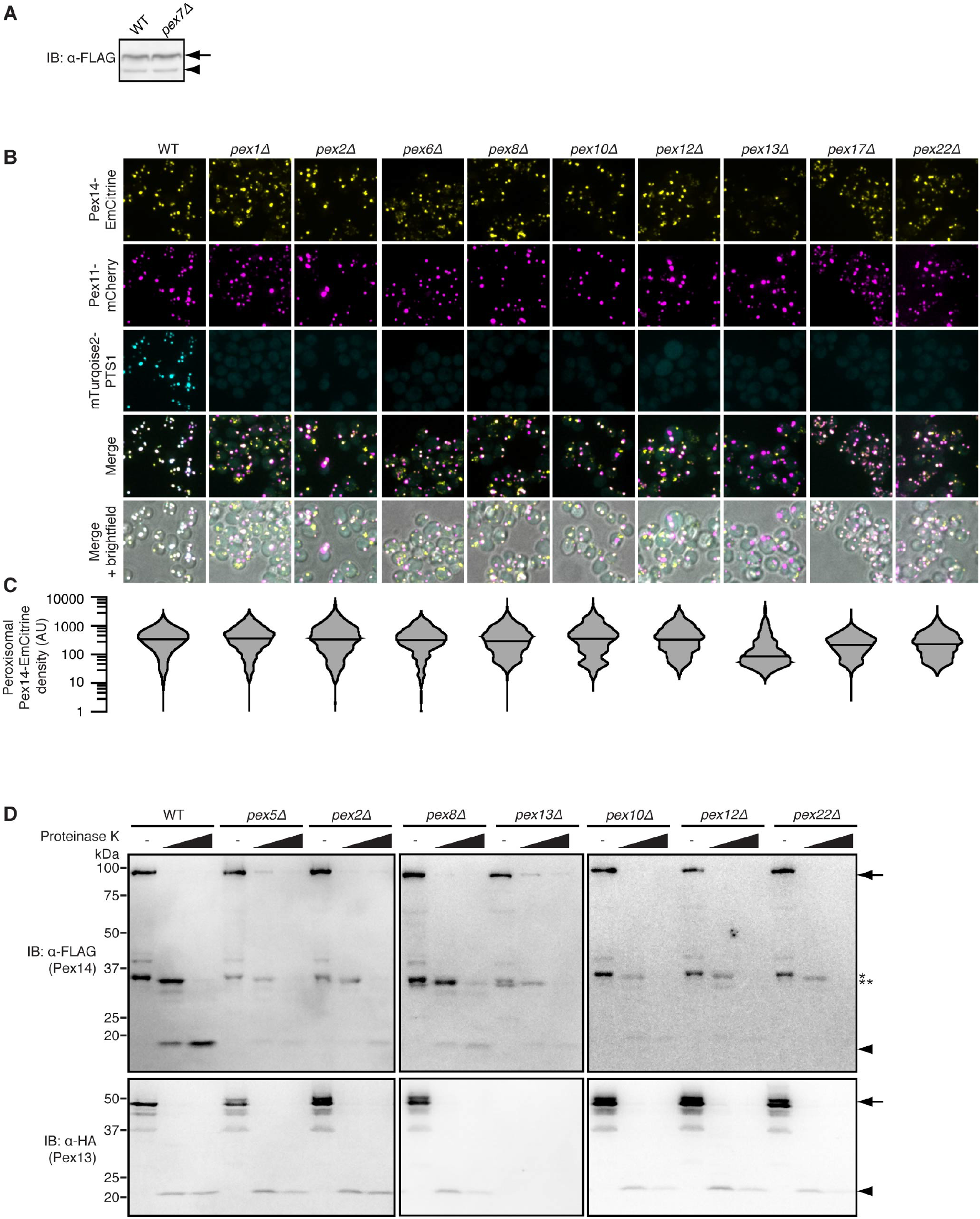
Additional evidence that Pex14 becomes arrested in a pre-integrated state on peroxisomes in importomer mutants. (A) Wild-type (WT) or *pex7Δ* cells expressing Pex14(hrv)-FLAG under the *PEX14* promoter from the *URA3* locus and cytosolic HRV 3C protease under the *GAL1* promoter were grown logarithmically for 15 hours in 2%/2% galactose/raffinose followed by 2 hours in 2% glucose. Extracts were resolved by SDS-PAGE and analyzed by immunoblotting (IB) with an anti-FLAG antibody. Arrow, full length Pex14(hrv)-FLAG; arrowhead, cleaved Pex14(hrv)-FLAG. (B) Representative confocal micrographs (maximum intensity projections) of logarithmically growing cells expressing Pex14-EmCitrine and the peroxisomal membrane and matrix markers Pex11-mCherry and the mTurquoise2-PTS1, respectively. Images from WT strains represent the same experiment as images from WT strains shown in Figure 3C. (C) Distributions of Pex14-EmCitrine density at Pex11-mCherry-defined peroxisomes in the images shown representatively in (B). Medians are represented as horizontal lines. Violin plot tails were cut off at 1 arbitrary units (AUs) for clarity of presentation. The distribution from the WT strain is identical to that shown in Figure 3D. 2865, 1536, 1155, 621, 1295, 842, 913, 977, 2708 and 1120 peroxisomes were analyzed for WT, *pex1Δ, pex2Δ, pex6Δ, pex8Δ, pex10Δ, pex12Δ, pex13Δ, pex17Δ* and *pex22Δ* strains, respectively. (D) Cells expressing FLAG-Pex14-MYC and Pex13-HA were grown to post-log phase before being subjected to spheroplasting and gentle lysis. Following centrifugation of lysates, membranes were resuspended in buffer and treated with proteinase K (doses: 0, 31 and 125 ng/μL) for 5 minutes at 37°C as indicated. Following proteinase K quenching, samples were resolved by SDS-PAGE and analyzed by immunoblotting (IB) with the indicated antibodies. Arrow, full length proteins; arrowhead, protected fragments; asterisk, proteinase K-independent degradation product; double asterisk, proteinase K-dependent partially digested intermediate.

**Figure S2.**
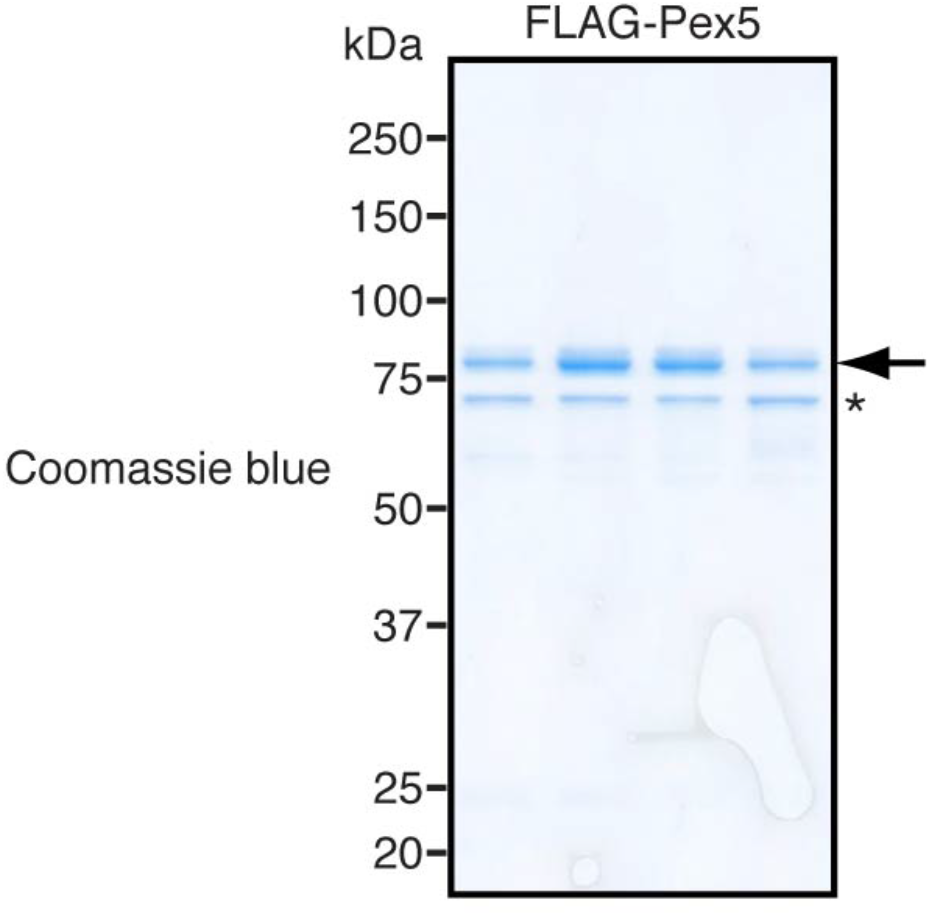
Purified recombinant FLAG-Pex5. FLAG-Pex5 gel filtration fractions were analyzed by SDS-PAGE and Coomassie blue staining. Pooled fractions are shown. Arrow, FLAG-Pex5; asterisk, probable FLAG-Pex5 degradation product.

**Figure S3.**
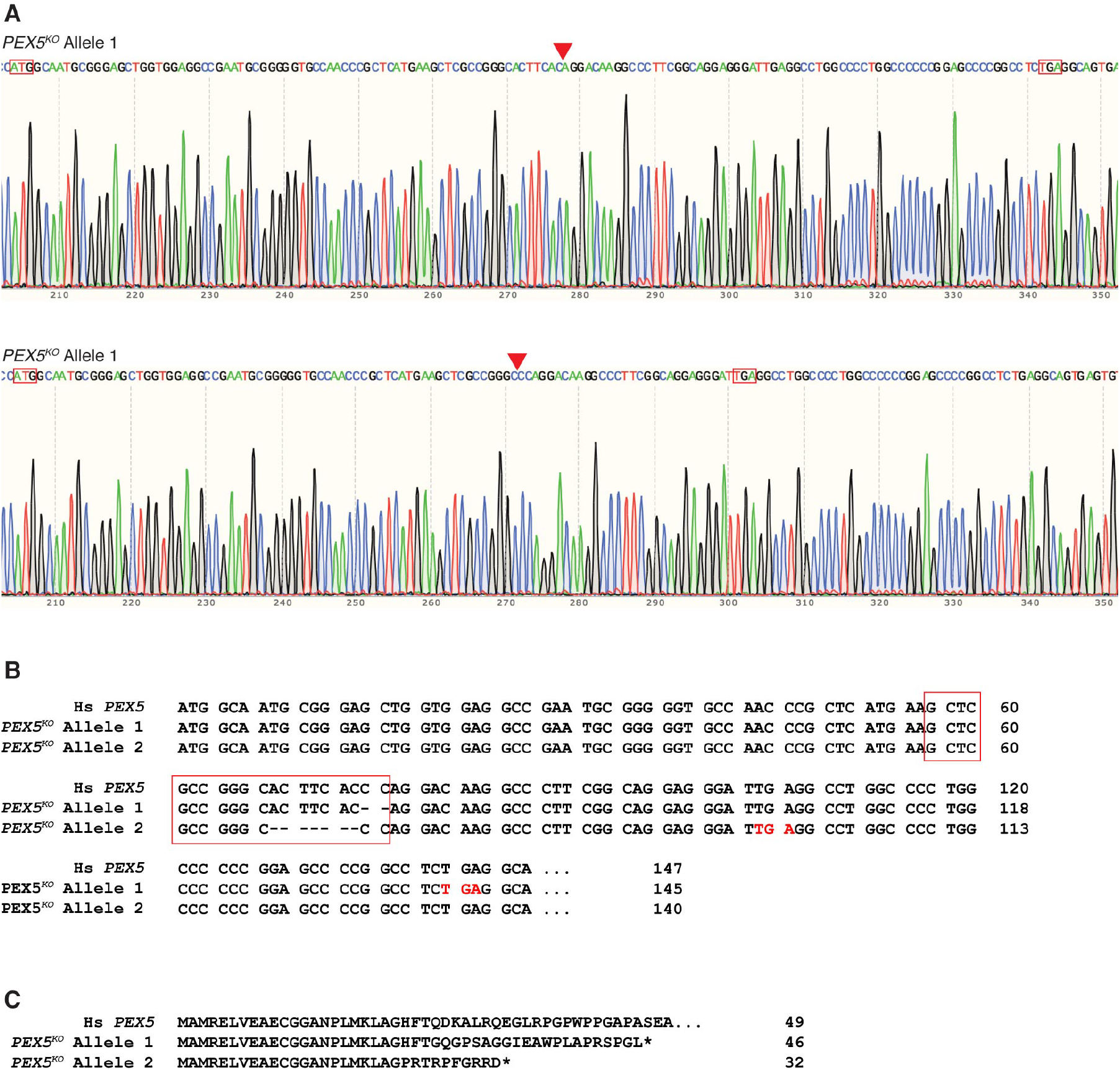
Sequence analysis of HEK293T *PEX5^KO^* alleles. (A) Chromatograms for *PEX5^KO^* allele sequencing. Start codons and premature stop codons are marked with red boxes. Deletion sites are marked by red arrows. (B) Nucleotide sequences of *PEX5^KO^* alleles. The WT allele (Hs *PEX5*) is shown for reference. The Cas9 target is marked with red boxes. Premature stop codons are highlighted in red. (C) Predicted amino acid sequences of *PEX5^KO^* allele translation products. The WT allele (Hs *PEX5*) is shown for reference. Asterisks indicate premature stop codons.

## MATERIALS AND METHODS

### Resource Table

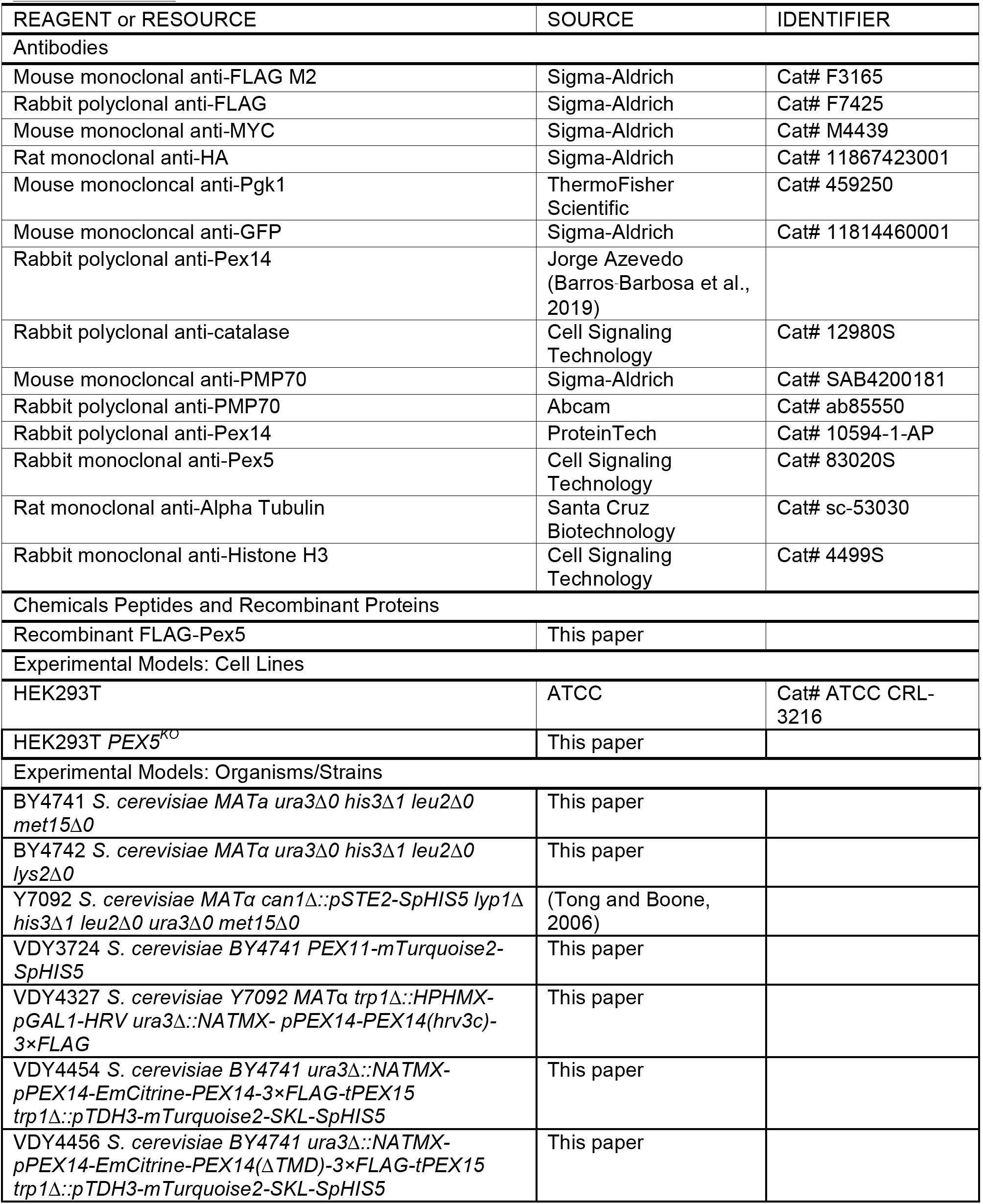
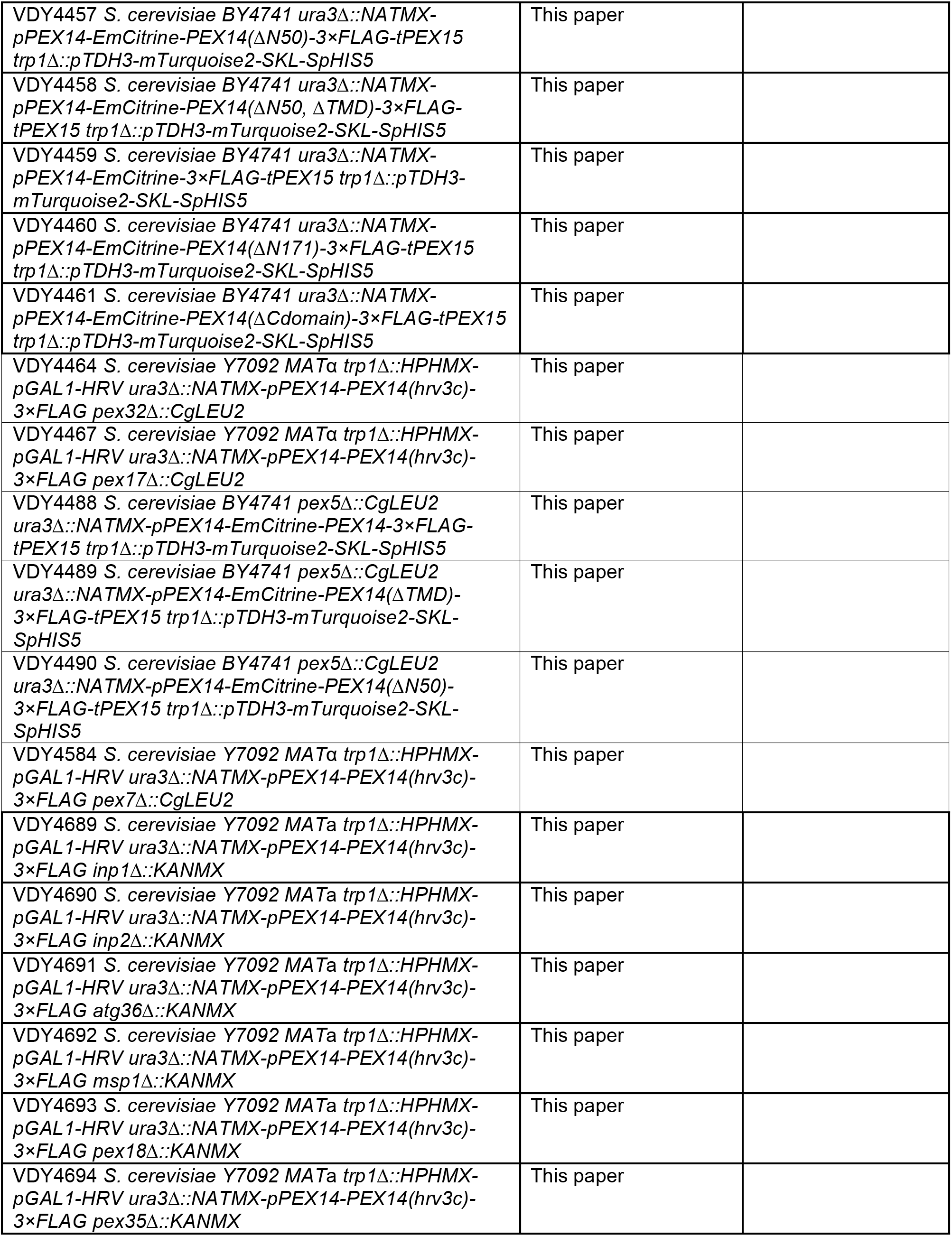
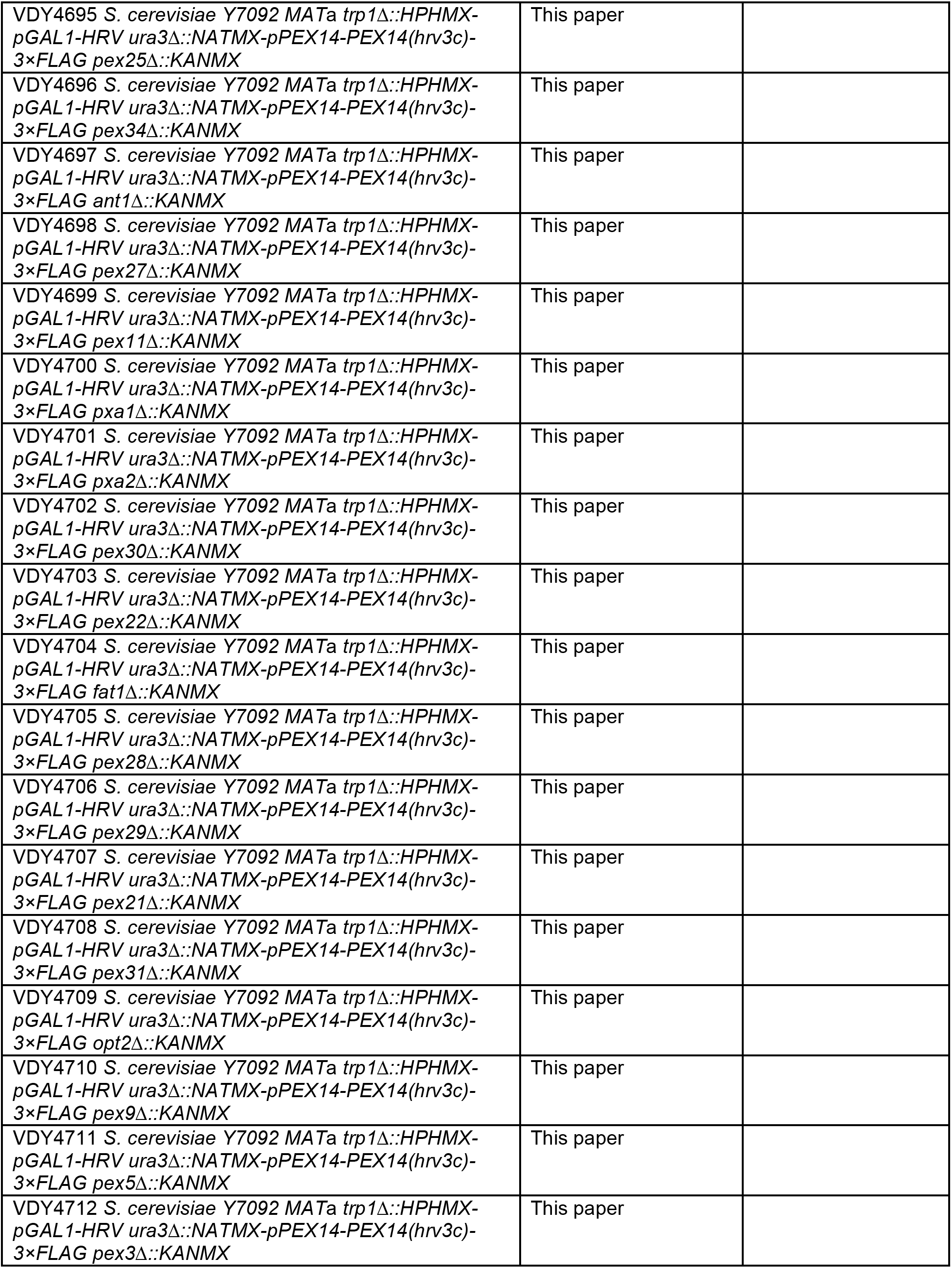
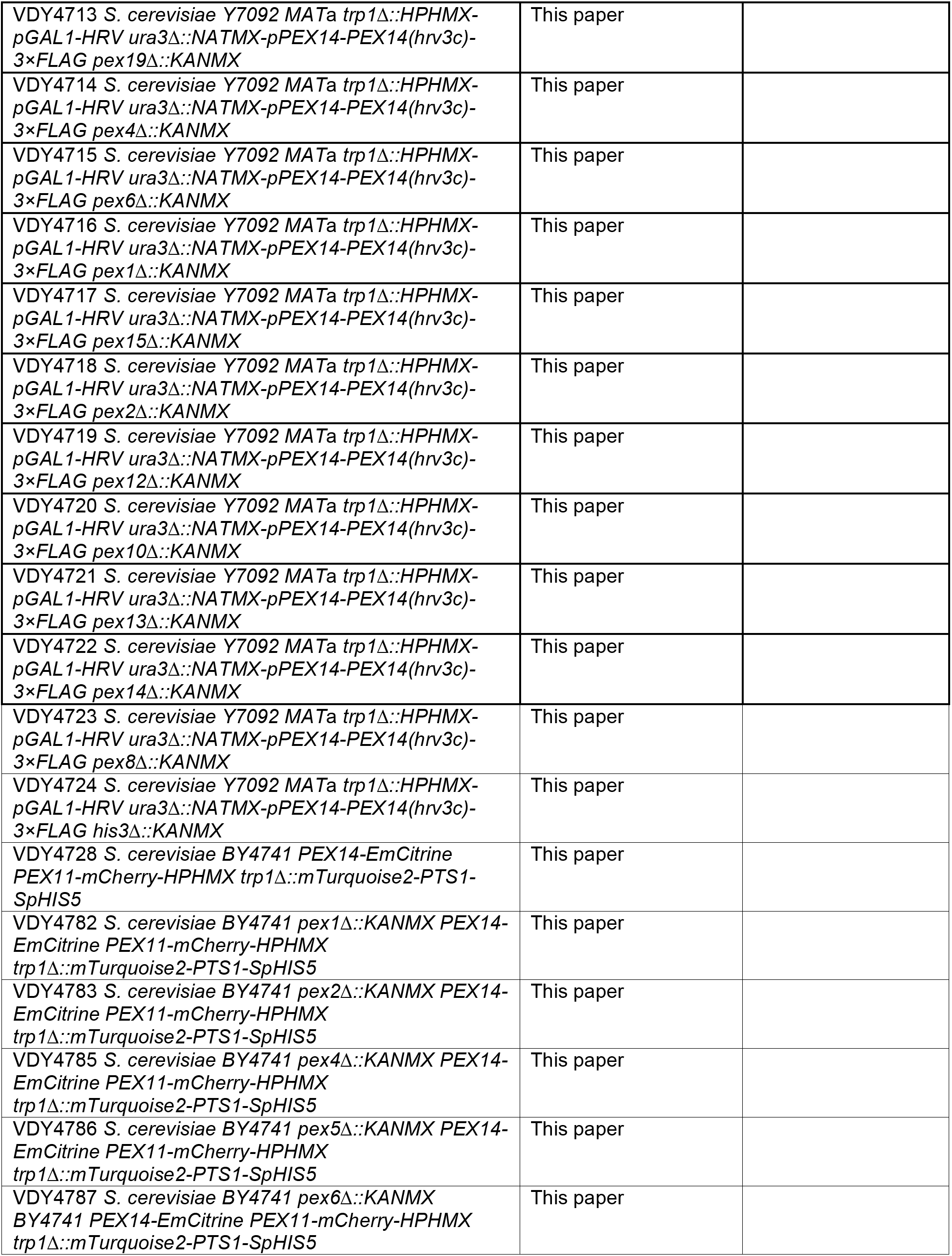
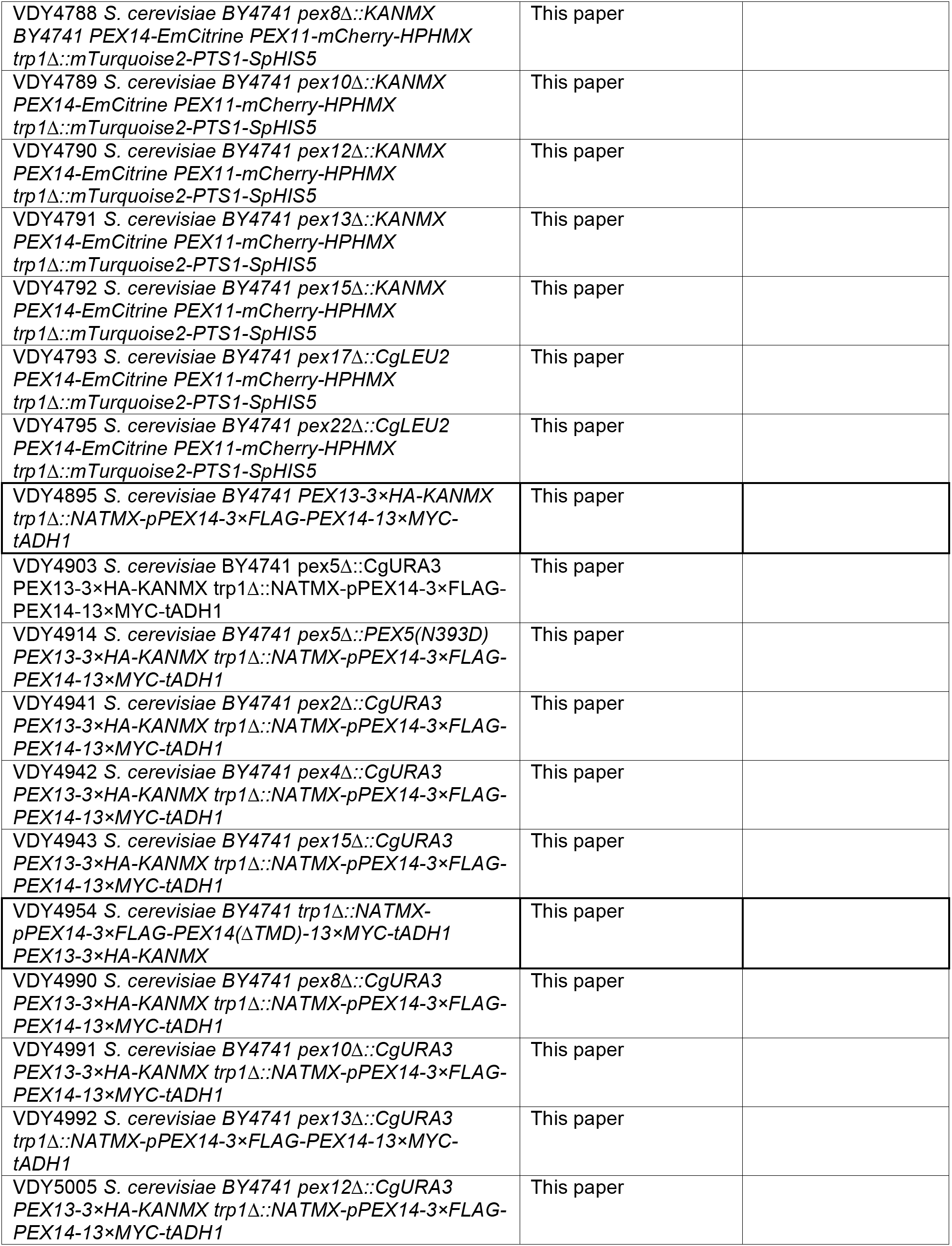
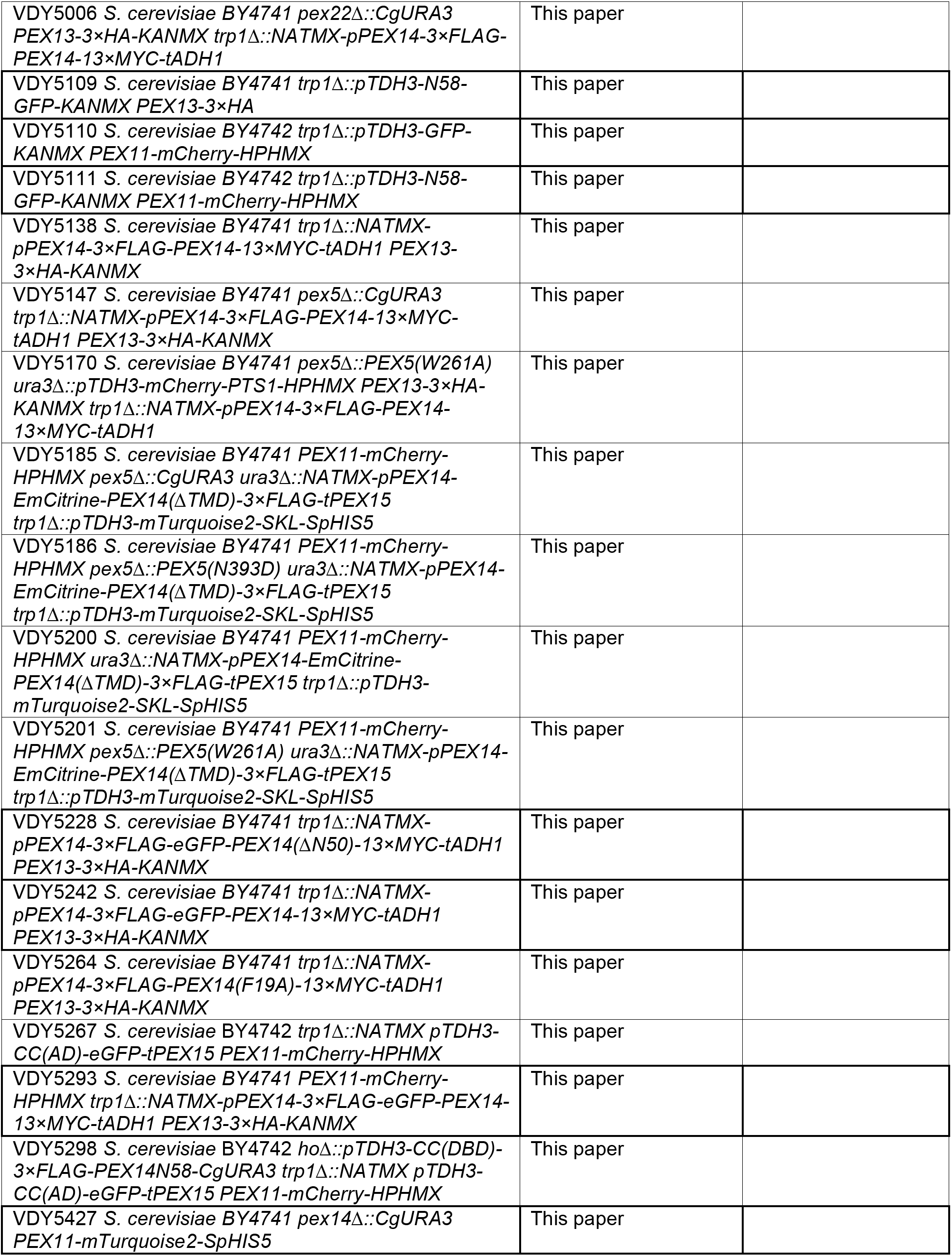
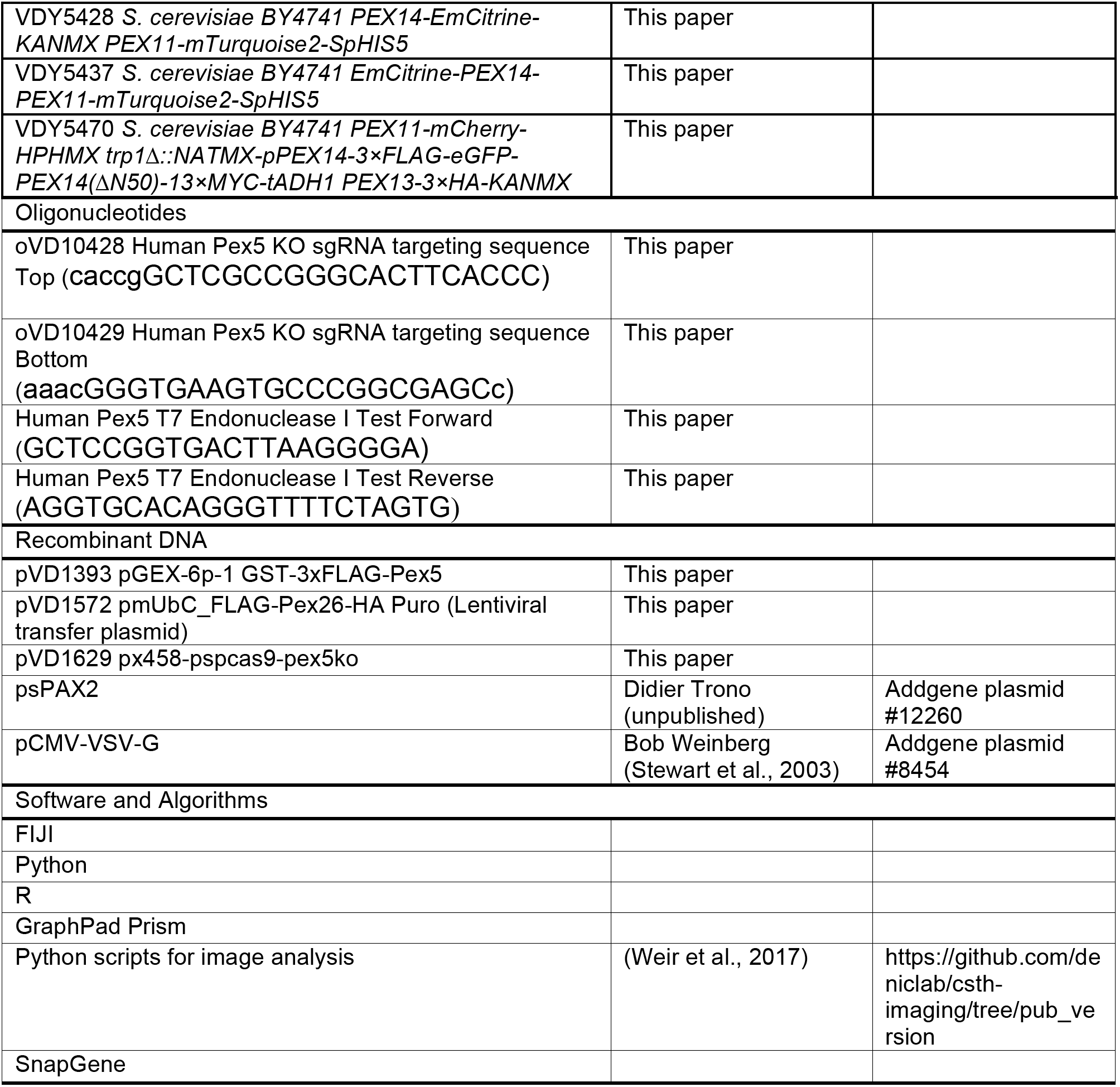

### Lead Contact and Materials/Data/Code Availability

Further information and requests for resources and reagents should be directed to and will be fulfilled by the Lead Contact, Vladimir Denic. All materials generated in this study, including strains, cell lines and plasmids, will be made available upon request. The authors declare that the data supporting the findings of this study are available within the paper and its supplementary information files. Python scripts for image analysis were previously reported (Weir et al., 2017) and are available at https://github.com/deniclab/csth-imaging/tree/pub_version.

### Experimental Model and Subject Details

*Saccharomyces cerevisiae* strains were derived from BY4741 (Mat **a**), BY4742 (Mat **α**) or Y7092 (MAT **α**) as indicated in the Resource Table. Liquid cultures were rotated continuously at 30°C. Cell growth on solid agar plates was carried out at 30°C followed by short-term storage at room temperature or under refrigeration. Strains were maintained as frozen glycerol stocks (−80°C) for long-term storage.

HEK293T cells were cultured in DMEM containing 10% FBS, 4 mM L-glutamine, 4.5 g/L glucose, 1 mM sodium pyruvate, 1.5 g/L sodium bicarbonate, 100 IU penicillin, and 100 μg/mL streptomycin at 37°C and 5% CO_2_. Blasticidin was added to 5 μg/mL to cells expressing lentiviral constructs. Cells were tested periodically for mycoplasma contamination using the LookOut Mycoplasma PCR Detection Kit from Millipore Sigma.

### Method Details

#### Yeast strain construction

All yeast strains were derived from *Saccharomyces cerevisiae* BY4741, BY4742 or Y7092. Gene deletions and gene expression cassette integrations were performed using standard PCR-based recombineering. Pex13 and Pex14 were fused to C-terminal 3×HA and EmCitrine, respectively, at their endogenous loci by standard PCR-based recombineering using tagging cassettes. Pex14 was fused seamlessly to an N-terminal EmCitrine by a standard procedure involving *URA3* knockin followed by 5-FOA counterselection. A similar procedure was used to make seamless allelic replacements at Pex14 and Pex5 endogenous loci. The HRV 3C protease screen library was constructed by crossing the query strain VDY4327 (derived from strain Y7092) to a homemade PMP gene deletion collection using synthetic genetic array methodology (Tong and Boone, 2006).

#### Yeast protease protection assay

Saturated yeast cultures were back diluted in YPD media (1% yeast extract, 2% peptone, 2% dextrose) to an OD_600_ of 0.02 and grown 15.5 hr at 30°C. Cells were then collected by centrifugation, washed once with 4°C water, then resuspended in 30°C softening buffer (0.1 M Tris pH 9.4, 10 mM DTT) and gently shaken at 30°C for 20 min. Cells were then washed with 30°C zymolyase buffer (1.2 M sorbitol, 42 mM K_2_HPO_4_, 8 mM KH_2_PO_4_, pH 7.4), followed by spheroplasting for 90 min at 30°C in zymolyase buffer containing 0.43 mg/mL zymolyase 20T (Amsbio 120491-1). The resulting spheroplasts were washed once with 30°C zymolyase buffer, then resuspended in 4°C homogenization buffer (0.6 M sorbitol, 10 mM Tris pH 7.4, 1 mM EDTA, 1 mM PMSF, 2× Roche cOmplete protease inhibitor cocktail, 0.2% BSA) and gently lysed with 15 passes of a dounce tissue homogenizer (Wheaton 62400-642) at 4°C. Cell debris was pelleted at 1,500 × g for 5 min at 4°C and discarded. The resulting supernatant was spun at 25,000 × g for 15 min at 4°C to obtain a pellet fraction with peroxisome membranes. Membranes were washed in 4°C SEM (250 mM sucrose, 1 mM EDTA, 10 mM MOPS/KOH pH 7.2) and finally resuspended in 4°C SEM. Where indicated, membranes were incubated in 1% Triton X-100 for 30 min on ice followed by treatment with proteinase K (Roche 3115879001) at the indicated concentrations at 37°C for 5 min. To quench proteinase K activity, samples were then treated with PMSF (5 mM) at 4°C for 10 min, before being diluted with pre-boiling 5/4× SDS-PAGE loading buffer (62.5 mM Tris-Cl pH 6.8, 2.5% SDS, 0.125% bromophenol blue, 12.5% glycerol, 125 mM β-mercaptoethanol) to a final loading buffer concentration of 1x, and further boiled for 10 min. Samples were analyzed by SDS-PAGE and immunoblotting.

#### Fluorescence *in situ* hybridization

Fluorescence *in situ* hybridization was permormed as described previously (Raj et al., 2008). Saturated yeast cultures were back diluted into complete synthetic media (0.67% yeast nitrogen base, 2% dextrose) to an OD_600_ of 8× 10^−4^ and grown 16 hr at 30°C, then treated with 3.7% formaldehyde for 45 min at room temperature. Fixed cells were then collected by centrifugation, washed once with 4°C Buffer B (1.2M sorbitol, 0.1M K_2_HPO_4_), then resuspended in Buffer B plus 2.5mg/mL zymolyase 20T (US Biological 37340-57-1) and spheroplasted for 1 hr at 30°C. The resulting spheroplasts were washed with 4°C Buffer B and resuspended in 4°C 70% EtOH followed by incubation overnight at 4°C. Permeabilized spheroplasts were then collected and washed in wash buffer (10% formamide (Ambion AM9342), 2×SSC (Thermofisher AM9763)). Hybridization with CAL Fluor Red 610-conjugated anti-*PEX14* mRNA FISH probes tiling the length of the *PEX14* ORF (0.125 μM, Biosearch Technologies) was carried out overnight at 30°C in hybridization buffer (10% formamide, 10% dextran sulfate, 2× SSC). Labeled samples were diluted with wash buffer and incubated an additional 1 hr at 30°C before being pelleted and resuspended in 2×SCC for analysis by quantitative confocal microscopy.

#### Yeast confocal microscopy and image quantitation

Saturated yeast cultures were back diluted into YPD media (1% yeast extract, 2% peptone, 2% dextrose) and grown 5 hr at 30°C to mid-log phase. Cells were applied to concanavalin A (MP Biomedicals, Santa Ana, CA)-treated coverslips, overlaid with glass slides and imaged at 25°C on a TI microscope (Nikon, Tokyo, Japan) equipped with a CSU-10 spinning disk (Yokogawa, Tokyo, Japan), an ImagEM EM-CCD camera (Hamamatsu, Hamamatsu, Japan), and a 100 × 1.45 NA objective (Nikon) and controlled with MetaMorph imaging software (Molecular Devices, Sunnyvale, CA).

For quantitative imaging, cultures were back diluted into complete synthetic media (0.67% yeast nitrogen base, 2% dextrose) and grown 5 hr at 30°C to mid-log phase, applied to concanavalin A-treated 8-well plates (Lab-Tek) and overlaid with a 1% agarose pad made with complete synthetic media (or 2×SCC for FISH experiments). Confocal z-stacks were acquired with 0.2 μm step size for 6 μm. See Quantitation and Statistical Analysis for details on image quantitation.

#### HRV 3C protease assay

Saturated yeast cultures were back diluted in raffinose media (0.67% yeast nitrogen base, 2% raffinose) and grown 5 hr at 30°C to mid-log phase, then back diluted in raffinose/galactose media (0.67% yeast nitrogen base, 2% raffinose, 2% galactose) to an OD_600_ of 0.01 and grown 15 hr at 30°C. Glucose was then added to each culture to a final concentration of 2%, and cultures were grown for an additional 2 hrs at 30°C prior to protein extraction.

#### Yeast protein extraction

Cells were pelleted and resuspended in 4°C NaOH (0.2 M) and incubated on ice for 10 min. Cells were again pelleted, resuspended in SDS-PAGE loading buffer (50 mM TrisC1 pH 6.8, 2% SDS, 0.1% bromophenol blue, 10% glycerol, 4% β-mercaptoethanol, 1 mM PMSF, 1x Roche cOmplete protease inhibitor cocktail) and boiled for 10 min. The resulting extracts were analyzed by SDS-PAGE and immunoblotting.

#### Recombinant FLAG-Pex5 expression and purification

BL21 (DE3)-RIPL competent Escherichia coli were transformed with plasmid pVD1393. A fresh colony was inoculated into 25 mL LB media (1% tryptone, 0.5% yeast extract, 0.5% NaCl) plus 100 mg/L ampicillin and grown overnight at 37°C to saturation, then back diluted into TB media (2% tryptone, 2.4% yeast extract, 0.4% glycerol, 0.017 M KH_2_PO_4_, 0.072 M K_2_HPO_4_) plus 100 mg/L ampicillin and grown at 37°C to an OD_600_ of 0.5. GST-FLAG-Pex5 expression was then induced with IPTG (1 mM) for 4 hr at 37°C. Cells were then pelleted, washed in PBS, flash-frozen in liquid nitrogen and stored at −80°C overnight. Frozen pellet was thawed at room temperature and resuspended in lysis buffer (25 mM HEPES-KOH pH 7.4, 100 mM NaCl, 100 mM KCl, 10% glycerol, 10 mM MgCl2) supplemented with 1x Roche cOmplete protease inhibitor cocktail and 1 mM PMSF and lysed via four 15,000 psi passes through an EmulsiFlex-C3 (Avestin, Inc). The resulting lysate was clarified at 16,000 × g at 4°C for 20 min and bound to a hand-packed glutathione sepharose 4B (GE17-0756-01) column. GST-FLAG-Pex5 was eluted at room temperature with lysis buffer supplemented with 20 mM reduced glutathione. This buffer was then exchanged for cleavage buffer (50 mM Tris-HCL, pH 6.8, 150 mM NaCl, 10% glycerol, 1 mM EDTA, 1 mM DTT) using a PD10 desalting column (GE Healthcare). The GST tag was removed via addition of Prescission protease (GE 27084301) at a concentration of 2 units per 100 μg recombinant protein, followed by incubation at 4°C for 15 hr. Prescission protease, any uncleaved GST-FLAG-Pex5 and free GST were removed via passage through a hand-packed glutathione sepharose 4B column. The flow through was concentrated using a spin concentrator followed by Superdex 200 10/300 GL gel filtration in lysis buffer.

#### *In vitro* translation and immunoprecipitation

Yeast translation extracts were prepared as described previously (Schuldiner et al., 2008). Saturated BY4741 yeast culture was back diluted into YPD and grown at 30°C to an OD_600_ of 2. Cells were pelleted, washed sequentially in water then buffer A (30 mM HEPES, 100 mM KOAc, 2 mM Mg(OAc)_2_, pH 7.4) supplemented with 2 mM DTT, resuspended in buffer A supplemented with 2 mM DTT, 2× Roche cOmplete protease inhibitor and 14% glycerol and flash frozen drop wise in liquid nitrogen. The sample was then ground in a Retsch PM100 ball mill and the resulting powder stored at −80°C overnight. Powder was nutated at 4°C until thawed, resuspended in buffer A supplemented with 2 mM DTT, 2×Roche cOmplete protease inhibitor and 14% glycerol and clarified by centrifuging at 12,000 × g at 4°C for 10 min. The supernatant was then centrifuged in an SW55-Ti swing bucket rotor at 29,000 × g 4°C for 30 min. The clear supernatant layer was collected and run over five serially connected Hi-Trap desalting columns (GE healthcare, 17-1408-01) in buffer A supplemented with 2 mM DTT and 14% glycerol. Fractions with OD_260_ > 50 were collected and pooled. Endogenous mRNA was removed by treating the pooled fractions (200 μL) with 2.4 μL CaCl2 (40 mM stock) and 1 μL micrococcal nuclease (New England BioLabs M0247S) at room temperature for 10 min. Samples were then quenched with 3.6 μL EGTA (100 mM stock) at room temperature for 5 min.

Capped mRNAs were *in vitro* transcribed from the appropriate PCR products using the mMESSAGE mMACHINE T7 kit as described previously (Schuldiner et al., 2008).

*In vitro* translation of ^35^S methionine-labeled proteins in yeast translation extracts was performed as described previously (Schuldiner et al., 2008). 28.5 μL extracts were combined with 7.5 μL 6x translation buffer (132 mM HEPES-KOH pH 7.4, 720 mM KOAc, 9 mM Mg(OAc)_2_, 4.5 mM rATP (Promega E601B), 0.6 mM rGTP (Promega E603B), 150 mM creatine phosphate (Roche 10621714001), 0.24 mM amino acid mix – met (Promega L9961), 10.2 mM DTT), 1.5 μL CPK mixture (10 mg/mL creatine Kinase (Roche 10127566001), 50% glycerol, 20 mM HEPES-KOH pH7.4), 1.5 μL RiboGuard RNAse inhibitor (Lucigen 75935-554), 3 μL ^35^S-labeled methionine (PerkinElmer NEG709A500UC), 2.5 μL recombinant protein (~450ng) and 300 ng capped mRNA and incubated at room temperature for 30 min. Further translation was arrested by addition of cycloheximide to a final concentration of 1 mM before subjecting the samples to immunoprecipitation analysis.

Anti-FLAG antibody (Sigma F3165) was pre-bound to protein G Dynabeads (Invitrogen 10004D) in buffer A supplemented with 14% glycerol. *In vitro* translation reactions were mixed with these anti-FLAG beads and shaken at 4°C for 1 hr. Beads were washed 2× with buffer A supplemented with 14% glycerol and boiled in SDS-PAGE loading buffer (50 mM Tris-Cl pH 6.8, 2% SDS, 0.1% bromophenol blue, 10% glycerol, 4% β-mercaptoethanol) for 10 min. Samples were resolved by SDS-PAGE. Association of FLAG-Pex5 with ^35^S methionine-labeled *in vitro* translation products was assessed by phosphorimage analysis using an Azure Sapphire imaging system.

#### Generation of HEK293T *PEX5^KO^* cell lines

Pex5 knock out cell lines were generated by homologous recombination using the CRISPR/Cas9 system. Briefly, sgRNA oligonucleotides targeting the human Pex5 locus (at 5’-GCTCGCCGGGCACTTCACCC-3’) were ligated into the BbsI site of pSpCas9(BB)-2A-GFP (PX458, Addgene Plasmid #48138) to generate pVD1629. HEK293T cells were transiently transfected with pVD1629 using Lipofectamine 3000 (Thermo Fisher Scientific). 24 hr post transfection, individual GFP-positive cells were sorted into 96-well plates using a BD Biosciences FACS Aria II cell sorter. Individual clones were verified by immunoblotting for Pex5 (Rabbit-anti-Pex5, 1:1000, Cell Signaling Technology) and sequencing of the Pex5 loci knockout alleles.

#### Generation of lentivirus

HEK293T cells were grown in Opti-MEM (Thermo Fisher Scientific) + 5% FBS for 48 hr to 90% confluency. Cells were then transfected via Lipofectamine 3000 (Thermo Fisher Scientific) with pVSV-G (Addgne Plasmid #8454, Stewart et al., 2003), pSPAX2 (Addgene Plasmid #12260) and a transfer plasmid encoding FLAG-Pex26-HA under control of the UbC promoter at a ratio of 1:3:4. Transfection media was replaced with fresh Opti-MEM + 5% FBS 8 hr post transfection. Media containing virus was collected 24 and 48 hr post transfection and pooled. Virus was then filtered through a 0.4 μM filter and frozen to −80°C in single-use aliquots.

#### HEK293T lentiviral transduction

Cells were incubated overnight in culture media containing lentivirus, 8 μg/mL polybrene, and lacking penicillin and streptomycin. Media was exchanged the next day to media lacking virus or polybrene and cells were allowed to grow for 24 more hr. Blasticidin was then added to 5 μg/mL to select for transduced cells.

#### HEK293T protease protection assay

HEK293T cells were grown to confluency in one 15 cm plate per condition. All subsequent steps were performed at 4°C. Cells were rinsed twice in 20 mL PBS and scraped into 1 mL of SEM buffer (250 mM sucrose, 1 mM EDTA, 20 mM MOPS-KOH, pH 7.4). Cells were then lysed with 30 passes through a tissue homogenizer (VWR Cat. No. 89026-386), followed by pelleting nuclei and un-lysed cells for 3 min at 3,000×g to obtain a post-nuclear supernatant (PNS). PNS was subsequently spun for 15 min at 25,000 × g to obtain a peroxisome-enriched membrane pellet. Membrane pellets were gently resuspended in 100 μl SEM and protein content was measured using a Pierce BCA Protein Assay Kit (Thermo Fisher Scientific) to normalize total protein concentration across samples. Protease protection reactions were carried out in SEM buffer for 40 min on ice using 200 μg/mL Proteinase K and where indicated 1% Triton X-100. To quench proteinase K activity, samples were then treated with 0.5 mg/mL PMSF at 4°C for 10 min, before being diluted with pre-boiling SDS-PAGE loading buffer and further boiled for 10 min. Samples were analyzed by SDS-PAGE and immunoblotting.

#### Immunofluorescence microscopy

HEK293T cells were seeded on acid-washed No. 1.5 cover glass in media as indicated above. Following washing with PBS, glass-adhered cells were treated with 4% paraformaldehyde for 15 min, followed by 0.5% Triton X-100. Fixed and permeabilized cells were then blocked with 10% Normal Goat Serum (NGS) dissolved in PBS + 0.1% Tween-20 (PBST) before antibody staining. Cells were stained with primary antibodies for one hr and secondary antibodies for 30 min, both at room temperature. Primary antibodies (Rabbit-anti-Catalase, Cell Signaling Technology; Mouse-anti-PMP70, Millipore Sigma; Rabbit-anti-PMP70, Cell Signaling Technology; Mouse-anti-FLAG M2, MilliporeSigma; Rabbit-anti-Pex14, ProteinTech) and secondary antibodies (Goat-anti-Mouse-Alexa-488, Thermo Fisher Scientific; Goat-anti-Rabbit-Cy5, Thermo Fisher Scientific) were diluted 1:1000 in PBST + 10% NGS. After three final PBST rinses, cover glass was mounted using ProLong Diamond + DAPI (Thermo Fisher Scientific) and allowed to cure for 48 hr. Slides were imaged on a Zeiss LSM 700 laser scanning confocal microscope (Harvard Center for Biological Imaging) equipped with 405 nm, 488 nm, and 638 nm lasers and a Variable Secondary Dichroic.

#### Quantification and Data Replication

Confocal microscopy images were normalized to background camera noise, peroxisomal or *PEX14* mRNA punctate structures were segmented, and the indicated fluorescence intensities within these segments were analyzed, using Python scripts developed previously (Weir et al., 2017). Data were then analyzed using R, and graphs were generated using GraphPad Prism. Sample sizes were chosen based on technically similar experiments published previously (Weir et al. 2017). Sequencing chromatograms were presented using SnapGene. Exact values of n and what n represents are indicated in figure legends. Findings in figures 1B, D-F; 2C-F; 3B-D; 4D, G, H; 5B, C; 6A-C; and S1A-C were successfully reproduced. Findings in figures 2B, G; 3E; 4C, E, F; S1D; S2; and S3 were deemed conclusive and thus no attempt was made to reproduce these data.

#### Data and Code Availability

This study did not generate any dataset/code.

